# Adaptation in protein fitness landscapes is facilitated by indirect paths

**DOI:** 10.1101/045096

**Authors:** Nicholas C. Wu, Lei Dai, C. Anders Olson, James O. Lloyd-Smith, Ren Sun

## Abstract

The structure of fitness landscapes is critical for understanding adaptive protein evolution (e.g. antimicrobial resistance, affinity maturation, etc.). Due to limited throughput in fitness measurements, previous empirical studies on fitness landscapes were confined to either the neighborhood around the wild type sequence, involving mostly single and double mutants, or a combinatorially complete subgraph involving only two amino acids at each site. In reality, however, the dimensionality of protein sequence space is higher (20^*L*^, *L* being the length of the relevant sequence) and there may be higher-order interactions among more than two sites. To study how these features impact the course of protein evolution, we experimentally characterized the fitness landscape of four sites in the IgG-binding domain of protein G, containing 20^4^ = 160,000 variants. We found that the fitness landscape was rugged and direct paths of adaptation were often constrained by pairwise epistasis. However, while direct paths were blocked by reciprocal sign epistasis, we found systematic evidence that such evolutionary traps could be circumvented by “extra-dimensional bypass”. Extra dimensions in sequence space – with a different amino acid at the site of interest or an additional interacting site – open up indirect paths of adaptation via gain and subsequent loss of mutations. These indirect paths alleviate the constraint on reaching high fitness genotypes via selectively accessible trajectories, suggesting that the heretofore neglected dimensions of sequence space may completely change our views on how proteins evolve.

---

The fitness landscape is a fundamental concept in evolutionary biology [1–6]. Large-scale datasets combined with quantitative analysis have successfully unraveled important features of empirical fitness landscapes [7–9]. Nevertheless, there is a huge gap between the limited throughput of fitness measurements (usually on the order of 10^2^ variants) and the vast size of sequence space. Recently, the bottleneck in experimen-tal throughput has been improved substantially by coupling saturation mutagenesis with deep sequencing [10–16], which opens up unprecedented opportunities to understand the structure of high-dimensional fit-ness landscapes [17–19].

Previous empirical studies on combinatorially complete fitness landscapes have been limited to subgraphs of the sequence space consisting of only two amino acids at each site (2^*L*^ genotypes) [20–25]. Adaptive walks in these subgraphs can only follow “direct paths”, where each mutational step reduces the Hamming distance from the starting point to the destination. In sequence space with higher dimensionality (20^*L*^, for a protein sequence with *L* amino acid residues), however, the extra dimensions may provide additional routes for adaptation. For example, some evolutionary dead ends (i.e. local maxima) may become sad-dle points and allow for further increase in fitness [26]. In this case, adaptation may proceed via “indirect paths” in sequence space, which involve extra mutations and reversions. The existence of indirect paths has been implied in different contexts [27, 28], but has not been studied systematically so its influence on protein adaptation remains unclear. Another underappreciated property of fitness landscapes is the influ-ence of higher-order interactions. Empirical evidence suggests that pairwise epistasis is prevalent in fitness landscapes [7, 22, 23, 29]. Specifically, sign epistasis between two loci is known to constrain adaptation by limiting the number of selectively accessible paths [20]. Higher-order epistasis (i.e. interactions among more than two loci) has received much less attention and its role in adaptation is yet to be elucidated [28,30].

In this study, we investigated the fitness landscape of all variants (20^4^ = 160,000) at four amino acid sites (V39, D40, G41 and V54) in an epistatic region of protein G domain B1 (GB1, 56 amino acids in total) (Supplementary Fig. 1), an immunoglobulin-binding protein expressed in Streptococcal bacteria [31, 32]. The four chosen sites contain 12 of the top 20 positively epistatic interactions among all pairwise interactions in protein GB1, as we previously characterized [33] (Supplementary Fig. 2). Thus the sequence space is expected to cover highly beneficial variants, which presents an ideal scenario for studying adaptive evolution. Briefly, a mutant library containing all amino acid combinations at these four sites was generated by codon randomization. The “fitness” of protein GB1 variants, as determined by both stability (i.e. the fraction of folded proteins) and function (i.e. binding affinity to IgG-Fc), was measured in a high-throughput manner by coupling mRNA display with Illumina sequencing (Methods, Supplementary Fig. 3A) [34,35]. The relative frequency of mutant sequences before and after selection allowed us to compute the fitness of each variant relative to the wild type protein (WT).

To understand the impact of epistasis on protein adaptation, we first analyzed subgraphs of sequence space including only two amino acids at each site (Fig. 1A). Each subgraph represented a classical adaptive landscape connecting WT to a beneficial quadruple mutant, analogous to previously studied protein fitness landscapes [9, 20]. Each variant is denoted by the single letter code of amino acids across sites 39, 40, 41 and 54 (for example, WT sequence is VDGV). Each subgraph is combinatorially complete with 2^4^ = 16 variants, including WT, the quadruple mutant, and all intermediate variants. We identified a total of 29 subgraphs in which the quadruple mutant was the only fitness peak. By focusing on these subgraphs, we essentially limited the analysis to direct paths of adaptation, where each step would reduce the Hamming distance from the starting point (WT) to the destination (quadruple mutant). Out of 24 possible direct paths, the number of selectively accessible paths (i.e. with monotonically increasing fitness) varied from 12 to 1 among the 29 subgraphs (Fig. 1B). In the most extreme case, only one path was accessible from WT to the quadruple mutant WLFA (Fig. 1A). We also observed a substantial skew in the computed probability of realization among accessible direct paths (Supplementary Fig. 4), suggesting that most of the realizations in adaptation were captured by a small fraction of possible trajectories [20]. These results indicated the ex-istence of sign epistasis and reciprocal sign epistasis, both of which may constrain the accessibility of direct paths [20, 36]. Indeed, we found that these two types of epistasis were prevalent in our fitness landscape (Fig. 1C). Furthermore, we classified the types of all 24 pairwise epistasis in each subgraph and computed the level of ruggedness as *f_sign_* + 2*f_reciprocal_*, where *f_type_* was the fraction of each type of pairwise epista-sis. As expected, the number of selectively inaccessible direct paths, i.e. paths that involve fitness declines, was found to be positively correlated with the ruggedness induced by pairwise epistasis (Fig. 1D, Pearson correlation = 0.66, p=1.0×10^−4^) [2].

**Figure 1.**
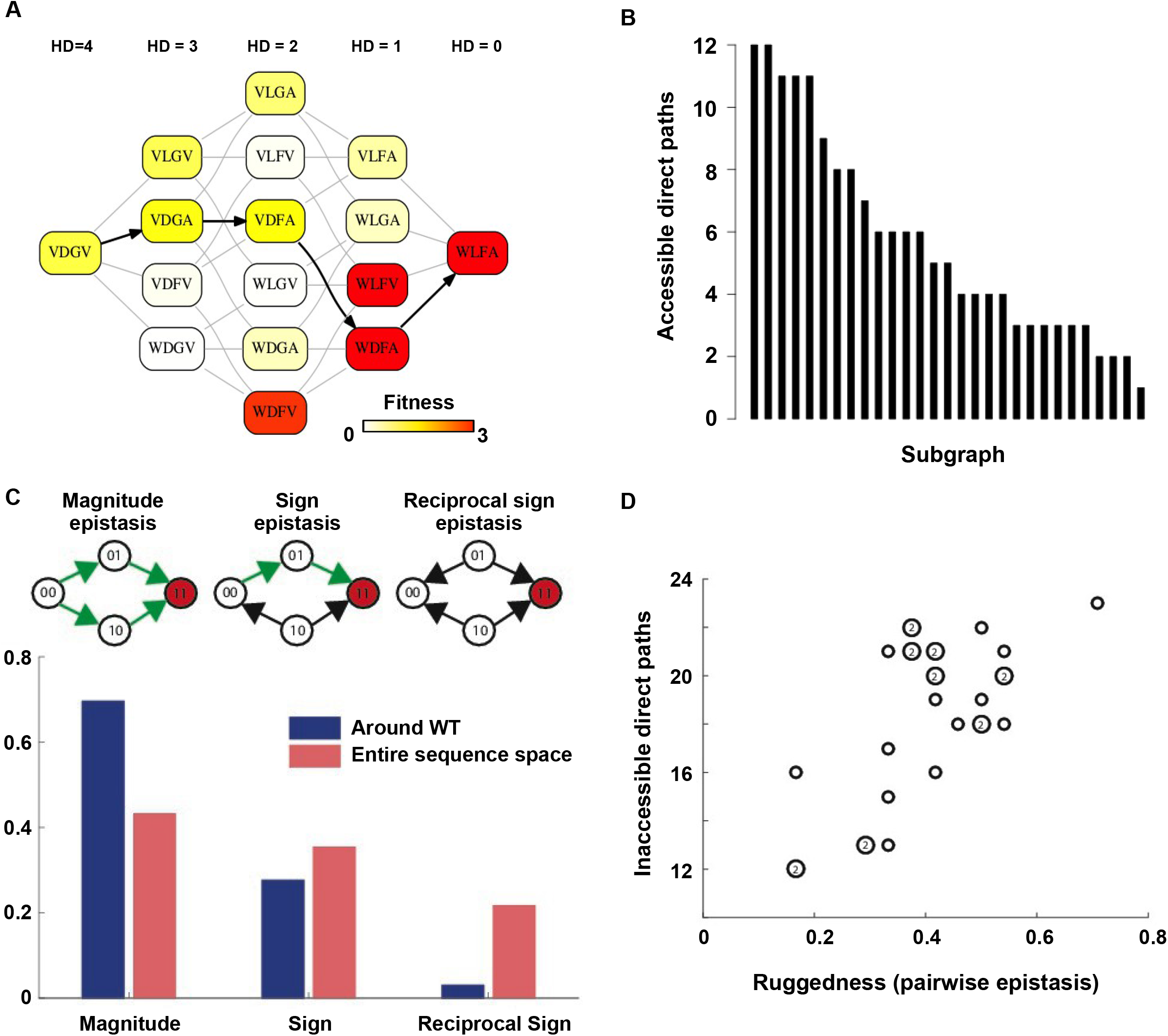
Direct paths of adaptation are constrained by pairwise epistasis. **(A)** An example of subgraph that contains VDGV (wild type, WT), the quadruple mutant WLFA and all intermediates between them. Each variant in the subgraph is represented by a node. Edges are drawn between nearest neighbors. The arrows in bold represent the only accessible direct path of adaptation from VDGV to WLFA. HD: Hamming Distance. **(B)** We identified a total of 29 subgraphs in which the quadruple mutant was the only fitness peak. The number of accessible direct paths from WT to the quadruple mutant is shown for each subgraph. The maximum number of direct paths is 24. **(C)** The fraction of three types of pairwise epistasis around WT (2091 out of 2166) or randomly sampled from the entire sequence space (10^5^ in total). Sign epistasis and reciprocal sign epistasis, both of which can block adaptive paths, are prevalent in the fitness landscape. Classification scheme of epistasis is shown at the top. Each node represents a genotype, which is within a sequence space of two loci and two alleles. Green arrows represent the accessible paths from genotype “00” to a beneficial double mutant “11” (colored in red). **(D)** The number of inaccessible direct paths are positively correlated (Pearson correlation = 0.66, p=1.0x10^−4^) with the ruggedness induced by sign and reciprocal sign epistasis. The level of ruggedness is quantified as *f_sign_* + 2*f*_*reciprocal*_, where *f_type_* denotes the fraction of each type of pairwise epistasis. The number inside a symbol indicates the number of subgraphs with identical properties.

Our findings support the view that direct paths of protein adaptation are often constrained by pairwise epistasis on a rugged fitness landscape [5,37]. In particular, adaptation can be trapped when direct paths are blocked by reciprocal sign epistasis. However, crucially, this analysis was limited to mutational trajectories within a subgraph of the sequence space. In reality, the dimensionality of protein sequence space is higher. Intuitively, when an extra dimension is introduced, a local maximum may become a saddle point and allow for further adaptation – a phenomenon recently proposed under the name “extra-dimensional bypass” [38]. We discovered two distinct mechanisms of bypass, either using an extra amino acid at the same site or using an additional site, that allow proteins to continue adaptation when no direct paths were accessible due to reciprocal sign epistasis (Fig. 2). The first mechanism of bypass, which we termed “conversion bypass”, works by converting to an extra amino acid at one of the interacting sites [28]. Consider a simple scenario with only two interacting sites. If the sequence space is limited to 2 amino acids at each site, as in past analyses of adaptive trajectories, the number of neighbors is 2; however, if all 20 possible amino acids were considered, the total number of neighbors would be 38. Some of these 36 extra neighbors may lead to potential routes that circumvent the reciprocal sign epistasis (Fig. 2A). In this case, a successful bypass would require a conversion step that substitutes one of the two interacting sites with an extra amino acid (00 → 20), followed by the loss of this mutation (21 → 11). This bypass is feasible only if the original reciprocal sign epistasis is changed to sign epistasis after the conversion. To test whether such bypasses were present in our system, we randomly sampled 10^5^ pairwise interactions from the sequence space and analyzed the ~20,000 reciprocal sign epistasis among them (Methods). More than 40% of the time there was at least one successful conversion bypass and in many cases multiple bypasses were available (Fig. 2B).

**Figure 2.**
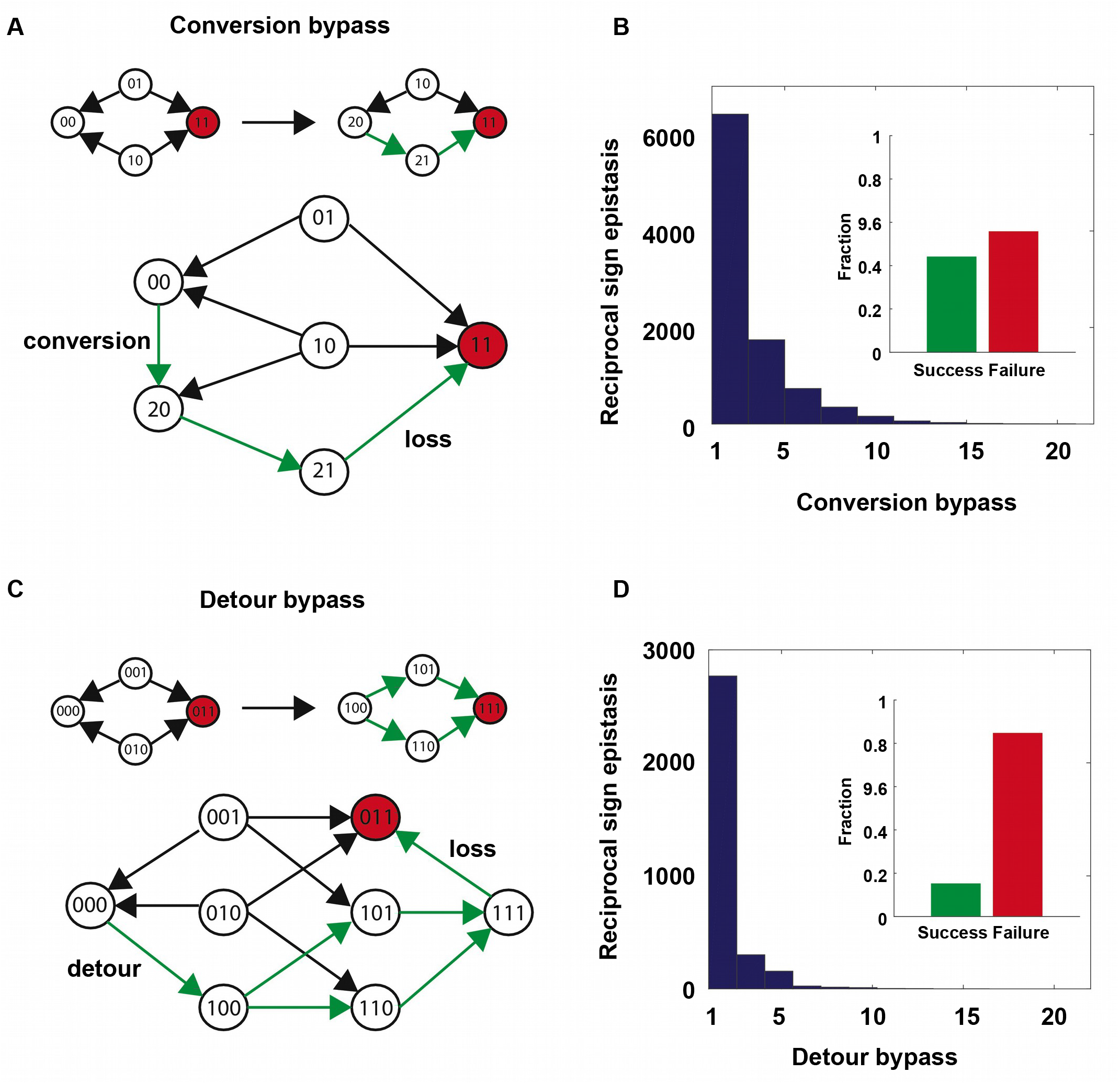
Two distinct mechanisms of extra-dimensional bypass. **(A)** Extra amino acids at one of the two interacting sites may open up potential paths that circumvent the reciprocal sign epistasis. The starting point is 00 and the destination is 11 (in red). Green arrows indicate the accessible path. A successful bypass would require a “conversion” step that substitutes one of the two interacting sites with an extra amino acid (00 → 20), followed by the loss of this mutation later (21 → 11). The original reciprocal sign epistasis is changed to sign epistasis on the new genetic background after conversion. **(B)** Among ~20,000 randomly sampled reciprocal sign epistasis, >40% of them can be circumvented by at least one conversion bypass (i.e. success, inset). The number of available bypass for the success cases is shown as histogram. **(C)** The second mechanism of bypass involves an additional site. In this case, adaptation involves a “detour” step to gain mutation at the third site (000 → 100), followed by the loss of this mutation (111 → 011). The original reciprocal sign epistasis is changed to either magnitude epistasis or sign epistasis on the new genetic background after detour (Supplementary Fig. 5). **(D)** In comparison to conversion bypass, detour bypass has a lower probability of success (<20%, inset) and is less prevalent.

The second mechanism of bypass, which we termed “detour bypass”, involves an additional site (Fig. 2C). In this case, adaptation can proceed by taking a detour step to gain a mutation at the third site (000 → 100), followed by the later loss of this mutation (111 → 011) [27,28]. Detour bypass was observed in our system (Fig. 2D), but was not as prevalent and had a lower probability of success than conversion bypass. Out of 38 possible detour bypasses for a chosen reciprocal sign epistasis, we found that there were on average 1.2 conversion bypasses and 0.27 detour bypasses available. We note, however, that the lower prevalence of detour bypass in our fitness landscape (*L*=4) does not necessarily mean that it should be expected to be less frequent than conversion bypass in other systems. While the maximum number of possible conversion bypasses is always fixed (19 × 2 – 2 = 36), the maximum number of possible detour bypasses (19 × (*L* – 2)) is proportional to the sequence length *L* of the entire protein (whereas our study uses a subset *L* = 4). The pervasiveness of extra-dimensional bypasses in our system contrasts with the prevailing view that adaptive evolution is often blocked by reciprocal sign epistasis, when only direct paths of adaptation are considered. The two distinct mechanisms of bypass both require the use of indirect paths, where the Hamming distance to the destination is either unchanged (conversion) or increased (detour).

In order to circumvent the inaccessible direct paths via extra dimensions, reciprocal sign epistasis must be changed into other types of pairwise epistasis. For detour bypass, this means that the original reciprocal sign epistasis is changed to either magnitude epistasis or sign epistasis in the presence of a third mutation (Supplementary Fig. 5A). There are three possible scenarios where detour bypass can occur (Supplementary Fig. 5B-D). We proved that higher-order epistasis is necessary for the scenario that reciprocal sign epistasis is changed to magnitude epistasis, as well as for one of the two scenarios that reciprocal sign epistasis is changed to sign epistasis (Supplementary Text). This suggests a critical role of higher-order epistasis in mediating detour bypass.

To confirm the presence of higher-order epistasis, we decomposed the fitness landscape by Fourier analysis (Fig. 3A, Methods) [9, 30]. The Fourier coefficients can be interpreted as epistatic interactions of different orders [6, 30], including the main effects of single mutations (the 1^*st*^ order), pairwise epistasis (the 2^*nd*^ order), and higher-order epistasis (the 3^*rd*^ and the 4^*th*^ order). The fitness of variants can be reconstructed by expansion of Fourier coefficients up to a certain order (Supplementary Fig. 6). In our system with four sites, the 4^*th*^ order Fourier expansion will always reproduce the measured fitness (i.e. Pearson correlation equals 1). When the 2^*nd*^ order Fourier expansion does not reproduce the measured fitness (i.e. Pearson cor-relation less than 1), it indicates the presence of higher-order epistasis. In this way, we identified the 0.1% of subgraphs with greatest fitness contribution from higher-order epistasis (Fig. 3A, red lines) and visual-ized the corresponding quadruple mutants by the sequence logo plot (Fig. 3B). The skewed composition of amino acids in these subgraphs indicates that higher-order interactions are enriched among specific amino acid combinations of site 39, 41 and 54. This interaction among 3 sites is consistent with our knowledge of the protein structure, where the side chains of sites 39, 41, and 54 can physically interact with each other at the core (Supplementary Fig. 1A) and destabilize the protein due to steric effects (Supplementary Fig. 7).

**Figure 3.**
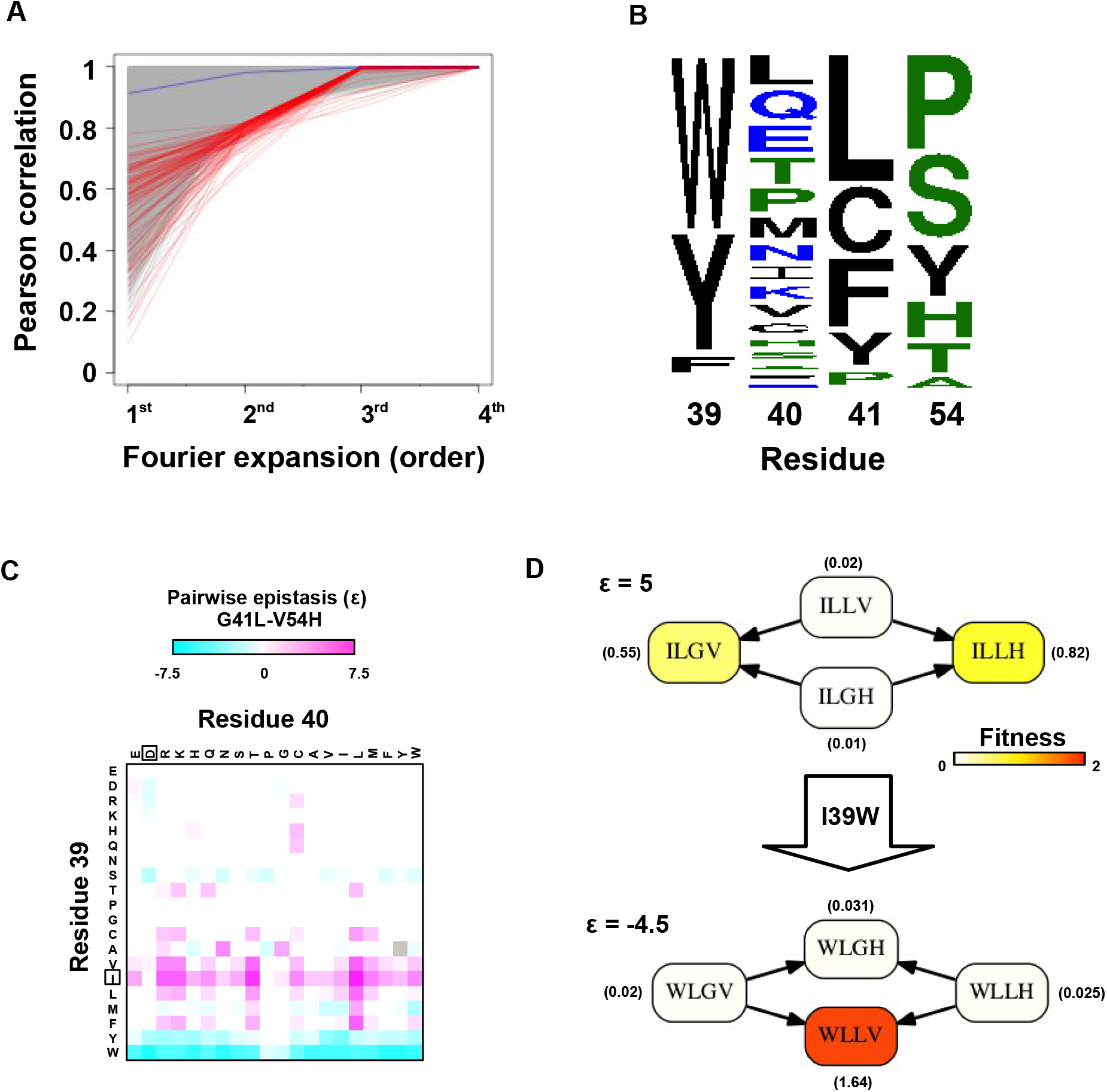
Evidence of higher-order epistasis. **(A)** The fitness decomposition was performed on all sub-graphs without missing variants. The fitness of variants can be reconstructed using Fourier coefficients truncated to a certain order. The Pearson correlation between the measured fitness and the fitness reconstructed by expansion of Fourier coefficients truncated to different orders (from 1^*st*^ to 4^*th*^) is shown for each subgraph. The blue line corresponds to the median Pearson correlation. The top 0.1% subgraphs with fitness contributions from higher-order epistasis (the bottom 0.1% subgraphs ranked by Pearson correlation at 2nd order expansion) are shown in red lines. **(B)** A sequence logo was generated for the quadruple mutants corresponding to the top 0.1% subgraphs with higher-order epistasis. The skewed composition of amino acids indicates that higher-order interactions are enriched among specific amino acid combinations of site 39, 41 and 54. **(C)** The magnitude of pairwise epistasis between G41L and V54H across different genetic backgrounds (i.e. all combinations of amino acids at site 39 and 40) is shown as a heat map. The amino acids of WT are boxed. Epistasis that cannot be determined due to missing variant is colored in grey. **(D)** Altering the genetic background at site 39 changed the positive epistasis (*ε* > 0) between G41L and V54H to negative epistasis (*ε* < 0). The fitness of each variant is indicated in the parentheses.

In the presence of higher-order epistasis, epistasis between any two sites would vary across different ge-netic backgrounds. We computed the magnitude of pairwise epistasis (ॉ) between each pair of amino acid substitutions (Methods)[39], and observed numerous instances where the sign of pairwise epistasis depended on genetic background. For example, G41L and V54H were positively epistatic when site 39 was isoleucine [I], but the interaction changed to negative epistasis when site 39 carried a tyrosine [Y] or a tryptophan [W] (Fig. 3C-D). Similar patterns were observed in other pairwise interactions among site 39, 41 and 54, such as G41F/V54A and V39W/V54H (Supplementary Fig. 8). The observed pattern of higher-order epistasis was consistent with the results of the Fourier analysis (Fig. 3B). For example, site 40 was mostly excluded from higher-order epistasis; tyrosine [Y] or tryptophan [W] at site 39 were involved in the most significant higher-order interactions, as they often changed the sign of pairwise epistasis. Higher-order epistasis can also switch the type of pairwise epistasis, such as shifting from reciprocal sign epistasis to magnitude or sign epistasis (Supplementary Fig. 9), which in turn is important for the existence of detour bypass.

Our analysis on circumventing reciprocal sign epistasis revealed how indirect paths could open up new avenues of adaptation. To study the impact of indirect paths at a global scale, we performed simulated adaptation in the entire sequence space of 160,000 variants. The fitness landscape was completed by im-puting fitness values of the 10,639 missing variants (i.e. 6.6% of the sequence space) that had fewer than 10 sequencing read counts in the input library. Our model of protein fitness incorporated main effects of single mutations, pairwise interactions, and three-way interactions among site 39, 41 and 54 (Methods, Supplementary Fig. 10). We used predictor selection based on biological knowledge, followed by regularized regression, which has been demonstrated to ameliorate possible bias in the inferred fitness landscape [40]. In the complete sequence space, we identified a total of 30 fitness peaks (i.e. local maxima); among them 15 peaks had fitness larger than WT and their combined basins of attraction covered 99% of the sequence space (Fig. 4A).

**Figure 4.**
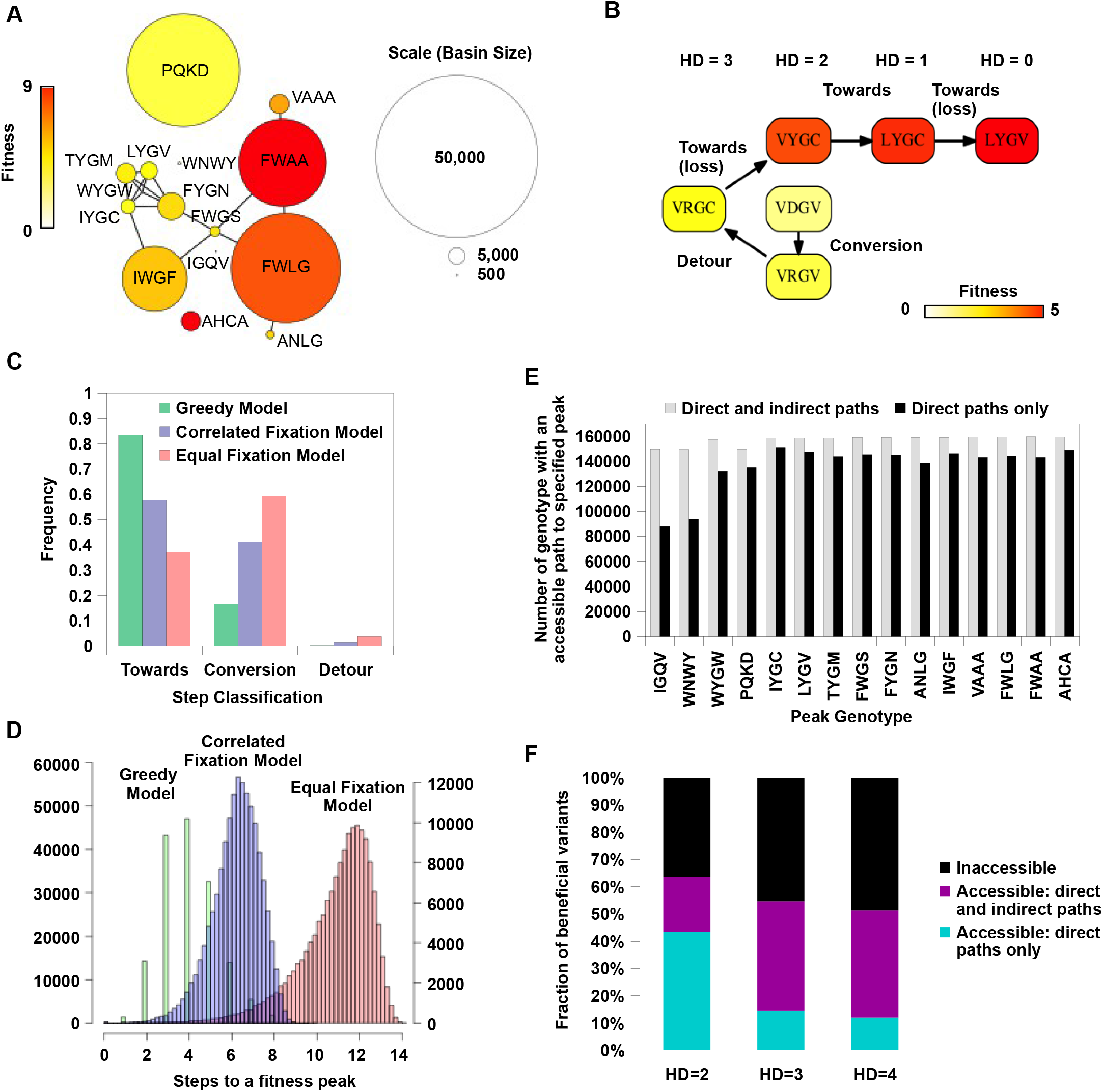
Indirect paths promote evolutionary accessibility. **(A)** 15 peaks had fitness larger than WT and their combined basins of attraction accounted for 99% of the entire sequence space. The size of each basin of attraction is identified by the Greedy Model (Methods). The area of each node is in proportion to the size of the basin of attraction of the corresponding fitness peak. An edge is drawn between fitness peaks that are separated by a Hamming distance of 2. **(B)** A possible adaptive path starting from WT (VDGV) to the fitness peak LYGV. **(C)** The frequency of different types of mutational step are shown. Three models, including the Greedy Model (green), Correlated Fixation Model (blue) and Equal Fixation Model (red), are used to simulate 1,000 adaptive paths starting from each variant in the sequence space. All the adaptive paths end at a fitness peak. **(D)** The distribution of the length of the adaptive path initiated at different starting points. For Correlated Fixation Model and Equal Fixation Model, the length was computed by averaging over 1,000 simulated paths for each starting point. The scale on the left is for Greedy Model. The scale on the right is for Correlated Fixation Model and Equal Fixation Model. **(E)** Indirect paths increased the number of genotypes accessible to each fitness peak. The 15 peaks are ordered by increasing fitness (from left to right). **(F)** A large fraction of beneficial variants in the sequence space (fitness > 1) were accessible from WT only via indirect paths.

We then simulated adaptation on the fitness landscape using three different models of adaptive walks (Meth-ods), namely the Greedy Model [6], Correlated Fixation Model [41], and Equal Fixation Model [20]. In the Greedy Model, adaptation proceeds by sequential fixation of mutations that render the largest fitness gain at each step. The other two models assign a nonzero fixation probability to all beneficial mutations, either weighted by (Correlated Fixation Model) or independent of (Equal Fixation Model) the relative fitness gain. Among all the possible adaptive paths to fitness peaks, many of them involved indirect paths, i.e. they em-ployed mechanisms of extra-dimensional bypass (Fig. 4B, Supplementary Fig. 11). We classified each step on the adaptive paths into three categories based on the change of Hamming distance to the destination (a fitness peak, in this case): “towards (-1)”, “conversion (0)”, and “detour (+1)” (Fig. 4C). Conversion was found to be pervasive during adaptation in our fitness landscape (17% of mutational steps for Greedy Model, 41% for Correlated Fixation Model, 59% for Equal Fixation Model). The use of detour was less frequent (0.1% of mutational steps for Greedy Model, 1.3% for Correlated Fixation Model, 3.7% for Equal Fixation Model), in accordance with the previous observation that detour bypass was less available than conversion bypass in our fitness landscape with *L* = 4. A conversion step would increase the length of an adaptive path by 1, while a detour step would increase the length by 2. As a result, an indirect path can be substantially longer than a direct path consisting of only “towards” steps. We found that many of the adaptive paths required more than 4 steps, which was the maximal length of a direct path between any variants in this landscape (Fig. 4D). Interestingly, because indirect adaptive paths involved more variants of intermediate fitness, the use of conversion and detour steps depended on the strength of selection. When mutations conferring larger fitness gains were more likely to fix (e.g. Greedy Model and Correlated Fixation Model), adaptation favored direct moves toward the destination, thus leading to a shorter adaptive paths (Fig. 4C-D). This suggests that the strength of selection interacts with the topological structure of fitness landscapes to determine the length and directness of evolutionary trajectories.

Given that extra-dimensional bypasses can help proteins avoid evolutionary traps, we expect that their existence would facilitate adaptation in rugged fitness landscapes. Indeed, we found that indirect paths increased the number of genotypes with access to each fitness peak (Fig. 4E). In addition, the fraction of genotypes with accessible paths to all 15 fitness peaks increased from from 34% to 93% when indirect adaptive paths were allowed (Supplementary Fig. 11C). We also found that a substantial fraction of beneficial variants (fitness > 1) in the sequence space were accessible from WT only if indirect paths were used (Fig. 4F). Taken together, these results suggest that indirect paths promote evolutionary accessibility in rugged fitness landscapes. This enhanced accessibility would allow proteins to explore more sequence space and lead to delayed commitment to evolutionary fates (i.e. fitness peaks) [28]. Consistent with this expectation, our simulations showed that many mutational trajectories involving extra-dimensional bypass did not fully commit to a fitness peak until the last two steps (Supplementary Fig. 12).

In our analysis, we have limited adaptation to the regime where fitness is monotonically increasing via sequential fixation of one-step beneficial mutants. When this assumption is relaxed, adaptation can some-times proceed by crossing fitness valleys [2, 6, 42, 43]. Another simplification in our analysis is to treat all sequences in a “protein space” [44], where two sequences are considered as neighbors if they differ by a single amino-acid substitution. In practice, amino acid substitutions occurring via a single nucelotide mutation are limited by the genetic code, so the total number of one-step neighbors would be reduced from 19*L* to approximately 6*L*. We also expect fitness landscapes of different systems to have different topo-logical structure. Even in our system (with >93% coverage of the genotype space), the global structure of the fitness landscape is influenced by the imputed fitness values of missing variants, which can vary when different fitness models or fitting methods are used. Our analysis also ignored measurement errors, but the measurement errors are expected to be very small due to the high reproducibility in the data (Supplementary Fig. 3B). Both imputation of missing variants and measurement errors can lead to slight mis-specification of the topological structure of the fitness landscape. Nevertheless, specific details of a certain fitness landscape do not undermine the generality of our findings on extra-dimensional bypass, higher-order epistasis, and their roles in protein evolution.

Higher-order epistasis has been reported in a few biological systems [28,45,46], and is likely to be common in nature [30]. In this study, we uncovered the presence of higher-order epistasis and systematically quanti-fied its contribution to protein fitness. We also revealed the importance of higher-order epistasis in mediating detour bypass, which could promote evolutionary accessibility in rugged fitness landscapes. As we pointed out, the possible number of detour bypasses scales up with sequence length, so it will be interesting to study how extra-dimensional bypass influences adaptation in sequence space of even higher dimensionality. For example, it is plausible that the sequence of a large protein may never be trapped in adaptation [47], so that adaptive accessibility becomes a quantitative rather than qualitative problem. Given the continuing develop-ment of sequencing technology, we anticipate that the scale of experimentally determined fitness landscapes will further increase, yet the full protein sequence space is too huge to be mapped exhaustively. Does this mean that we will never be able to understand the full complexity of fitness landscapes? Or perhaps big data from high-throughput measurements will guide us to find general rules? By coupling state-of-the-art experimental techniques with novel quantitative analysis of fitness landscapes, this work takes the optimistic view that we can push the boundary further and discover new mechanisms underlying evolution [9,48,49].

## Supplementary Figure Legends

**Supplementary Figure 1.**
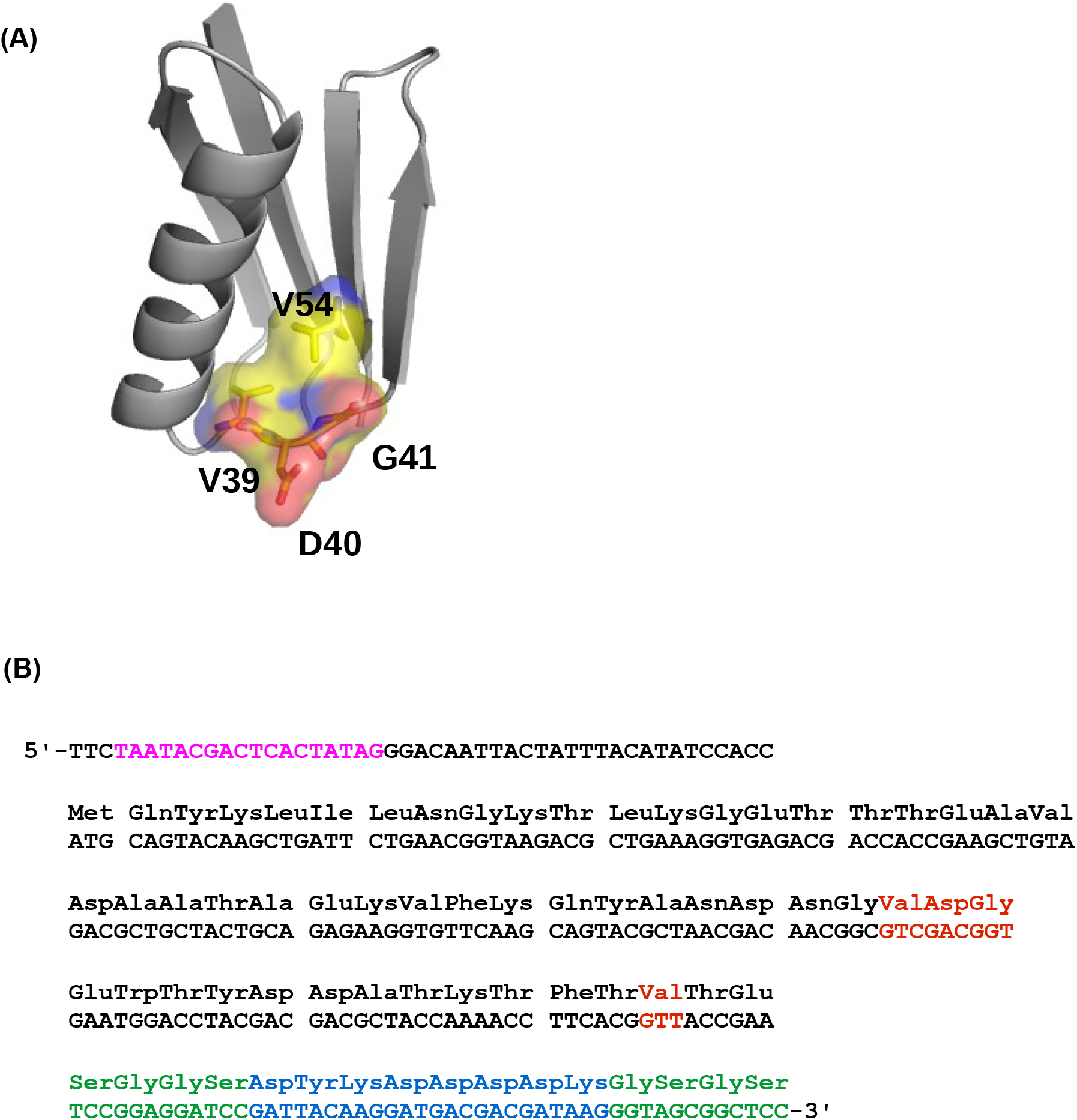
The four-site sequence space of protein G. **(A)** The locations of sites 39, 40, 41, and 54 of protein GB1 are shown on the protein structure. PDB: 1PGA [50]. **(B)** The WT sequence of the nucleotide template [33]. T7 promoter is highlighted in magenta. Randomized sites (39, 40, 41, and 54) are highlighted in red. Poly-GS linkers are highlighted in green. FLAG-tag is highlighted in blue.

**Supplementary Figure 2.**
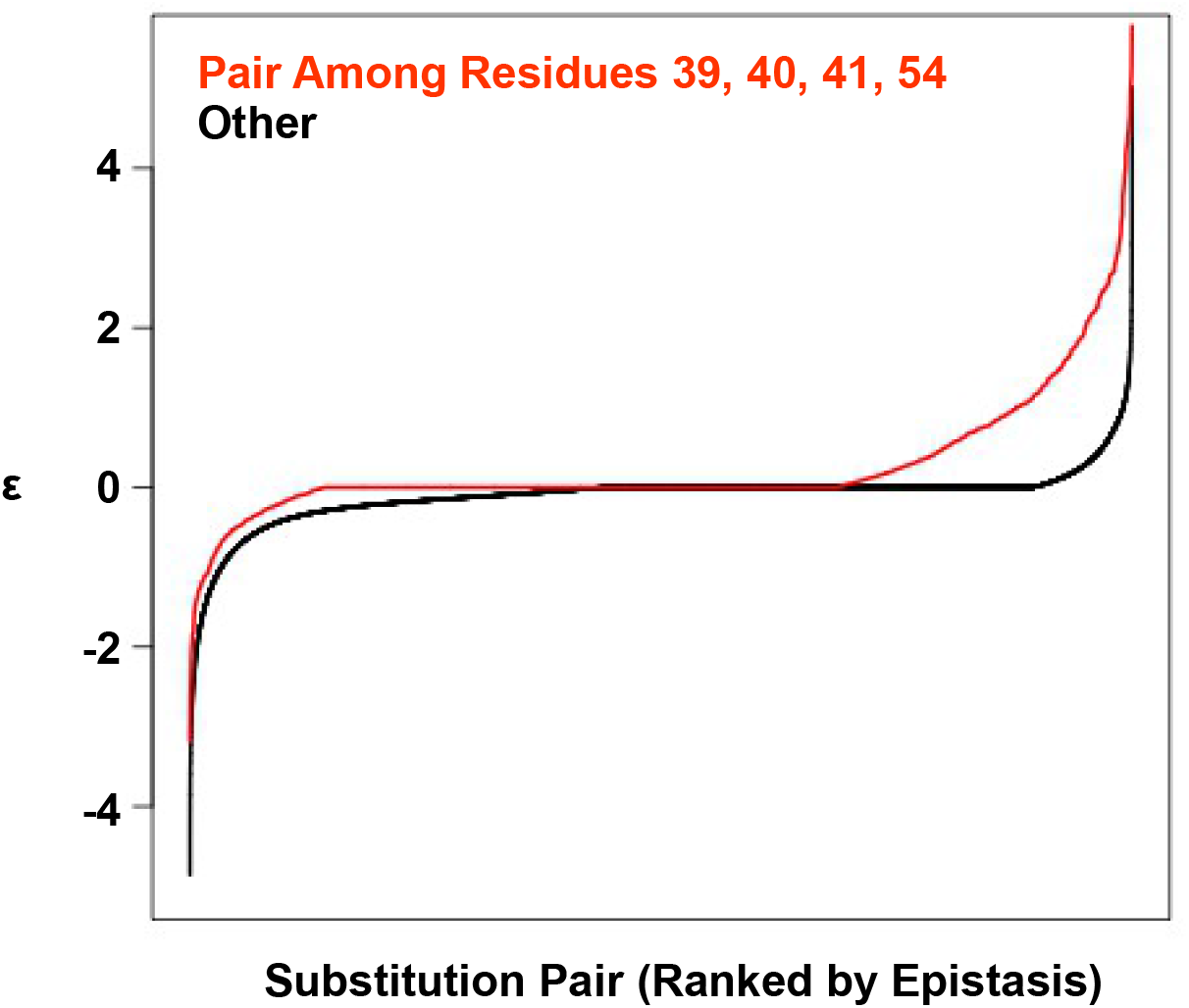
Positive epistasis is enriched in the four-site sequence space. The distribution of pairwise epistasis measured by Olson et al. [33]. The pairwise epistatic values among sites 39, 40, 41, and 54 are ranked and represented by the red line. The pairwise epistatic values among other sites (all but 39, 40, 41, and 54) are ranked and represented by the black line.

**Supplementary Figure 3.**
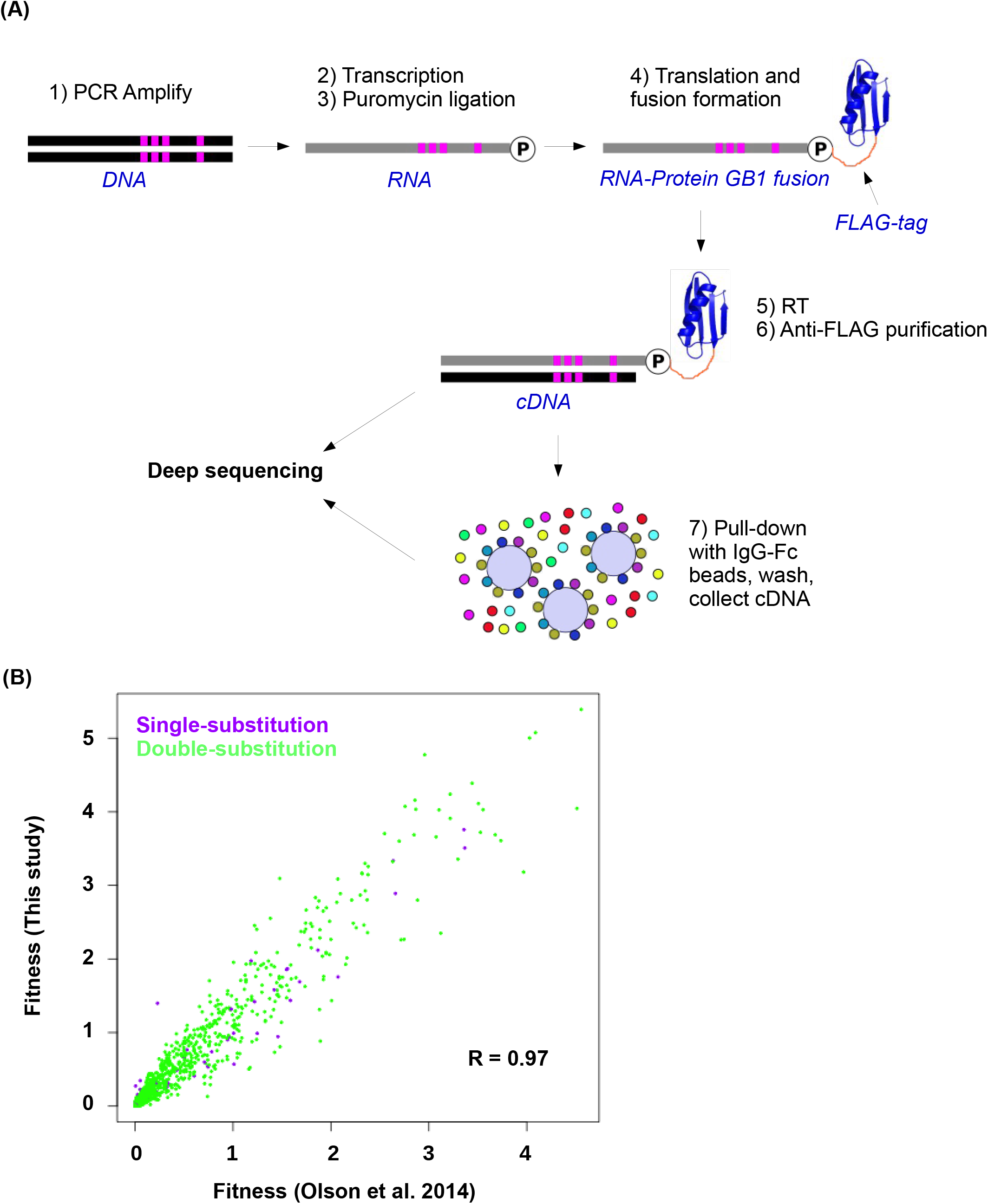
Workflow of mRNA display and data validation. **(A)** The workflow of mRNA display is shown. This is adapted from [33]. **(B)** The fitness values for all single substitution variants and double substitution variants in this study and in our previous study (based on an independently constructed library) [33] are compared. The high correlation (Pearson correlation=0.97) validates the fitness data ob-tained in this study.

**Supplementary Figure 4.**
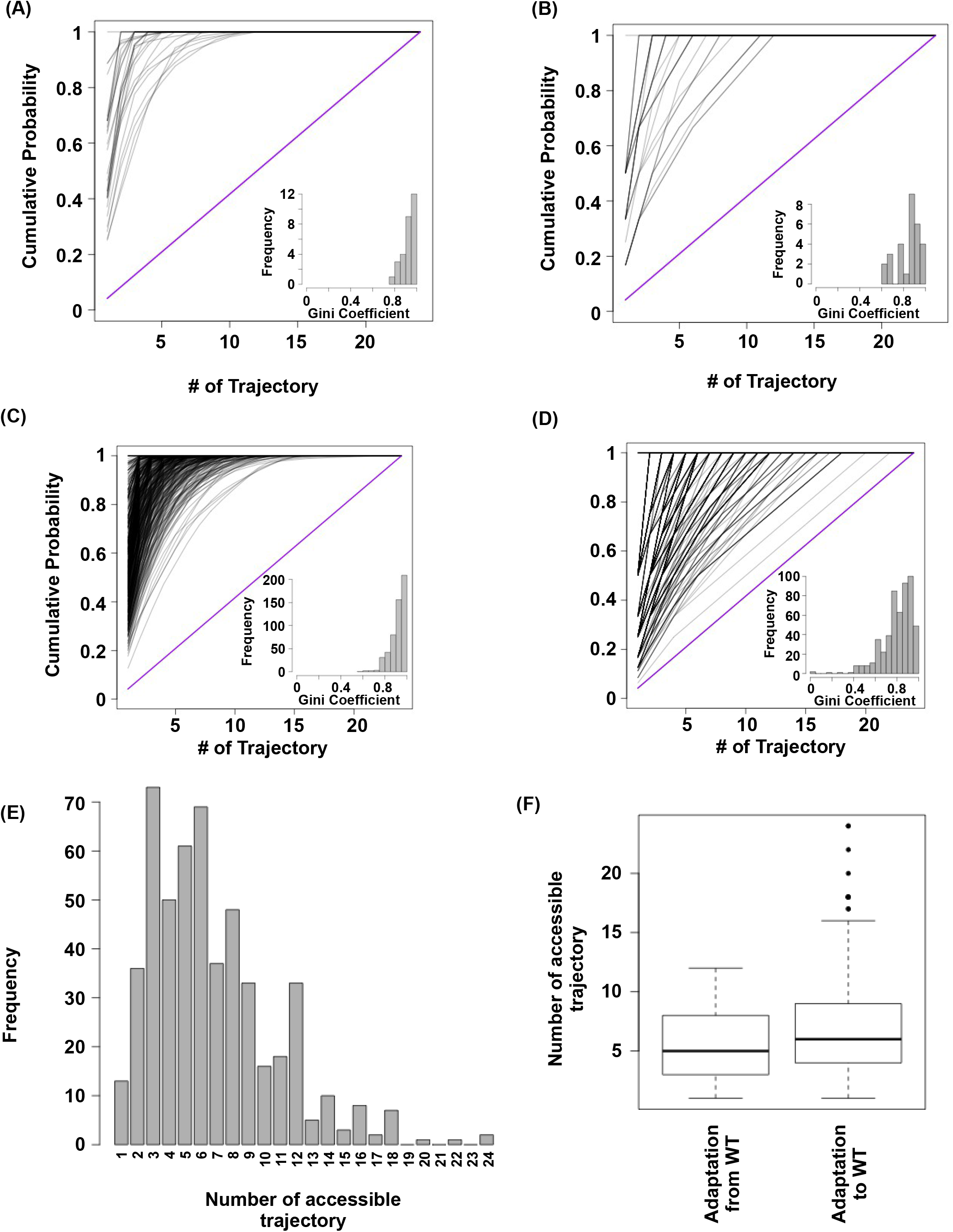
Subgraph analysis. We calculated the relative probabilities to realize each accessible path [20] (see Methods). In all the subgraphs analyzed, we found that most of the realizations were captured by a few accessible paths, as demonstrated by the skew in the cumulative probability of re-alization among different paths. **(A)** Cumulative probability of realization for mutational trajectories from WT (VDGV) to beneficial variants that have a Hamming distance (HD) of 4 from WT. This analysis only included those subgraphs with a reachable quadruple mutation variant (HD = 4 from WT) as the only fit-ness peak. Correlated Fixation Model is used. The diagonal line indicates the cumulative probability of a subgraph with equal probability of realization for all 24 possible trajectories. The bias of probability of realization in each subgraph was quantified using the Gini index (see Methods) and is shown as a histogram in the inset. **(B)** Same as panel A, except Equal Fixation Model is used instead. **(C and D)** Cumulative probability for mutational trajectories from a deleterious variants that have a Hamming distance (HD) of 4 from WT (VDGV) to WT. This analysis only included those subgraphs with WT being the only fitness peak and the quadruple variant has a fitness between 0.01 to 1. The diagonal line indicates the cumulative probability of a subgraph with equal probability of realization for all 24 possible trajectories. The bias of probability of realization in each subgraph was quantified using the Gini index and is shown as a histogram in the inset. A total of 526 subgraphs were analyzed. **(C)** Correlated Fixation Model is used. **(D)** Equal Fixation Model is used. **(E)** Number of accessible trajectory from a deleterious variants that have a Hamming distance (HD) of 4 from WT (VDGV) to WT in each subgraph is shown as a barplot. The maximum possible number of accessible trajectory is 24. **(F)** The distribution of number of accessible trajectory is shown as a box plot. “Adaptation from WT” indicates those subgraphs based on the adaptation from WT to a beneficial variant that has a Hamming distance of 4 from WT. “Adaptation to WT” indicates those sub-graphs based on the adaptation from a deleterious variant that has a Hamming distance of 4 from WT to WT.

**Supplementary Figure 5.**
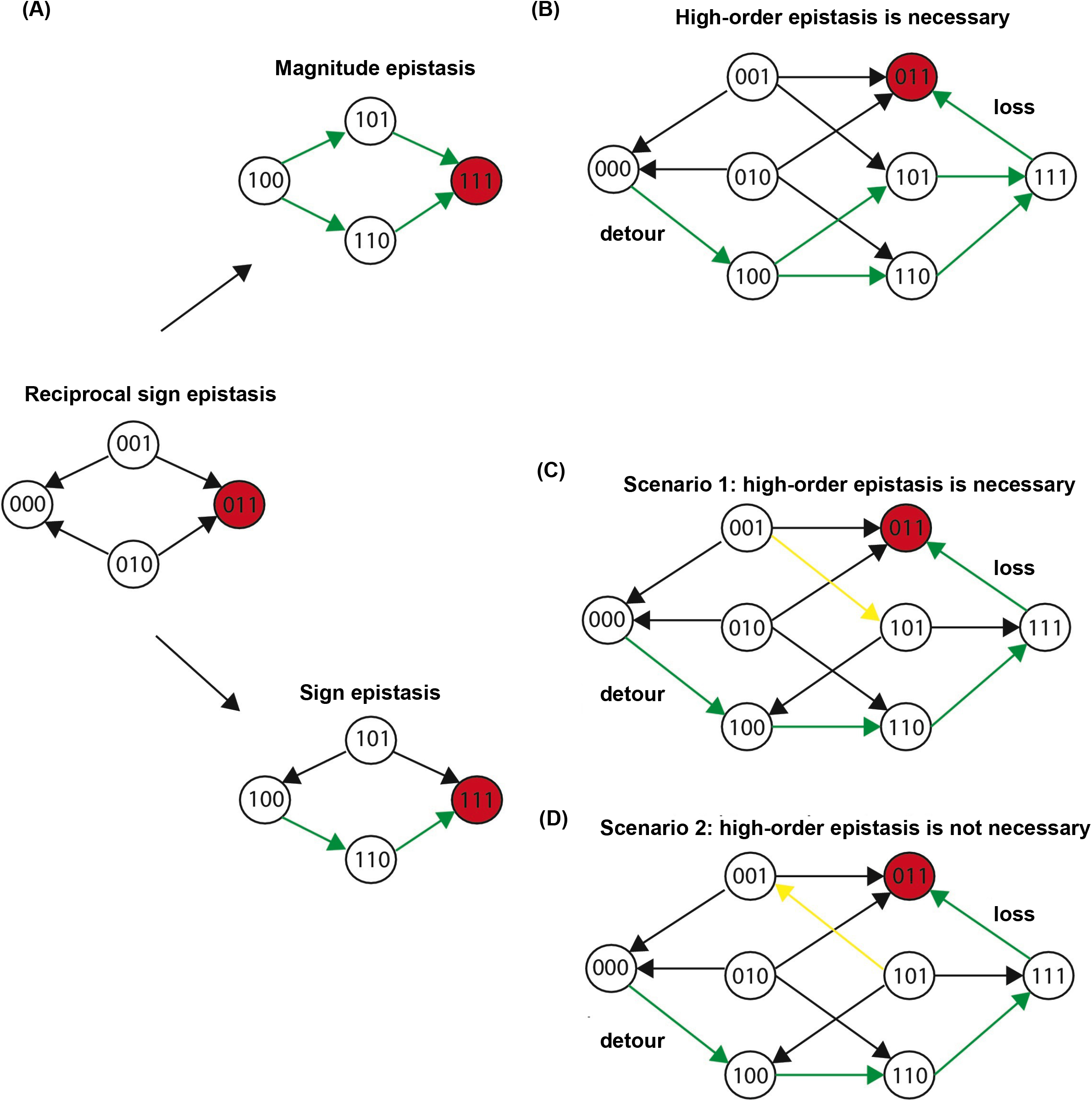
Three scenarios of extra-dimensional bypass via an extra site. **(A)** Reciprocal sign epistasis may be bypassed via the involvement of a third site. **(B-D)** There are three possible scenarios. It can be proven that higher-order epistasis is required for the scenarios in **(B)** and **(C)** (Supplementary Text).

**Supplementary Figure 6.**
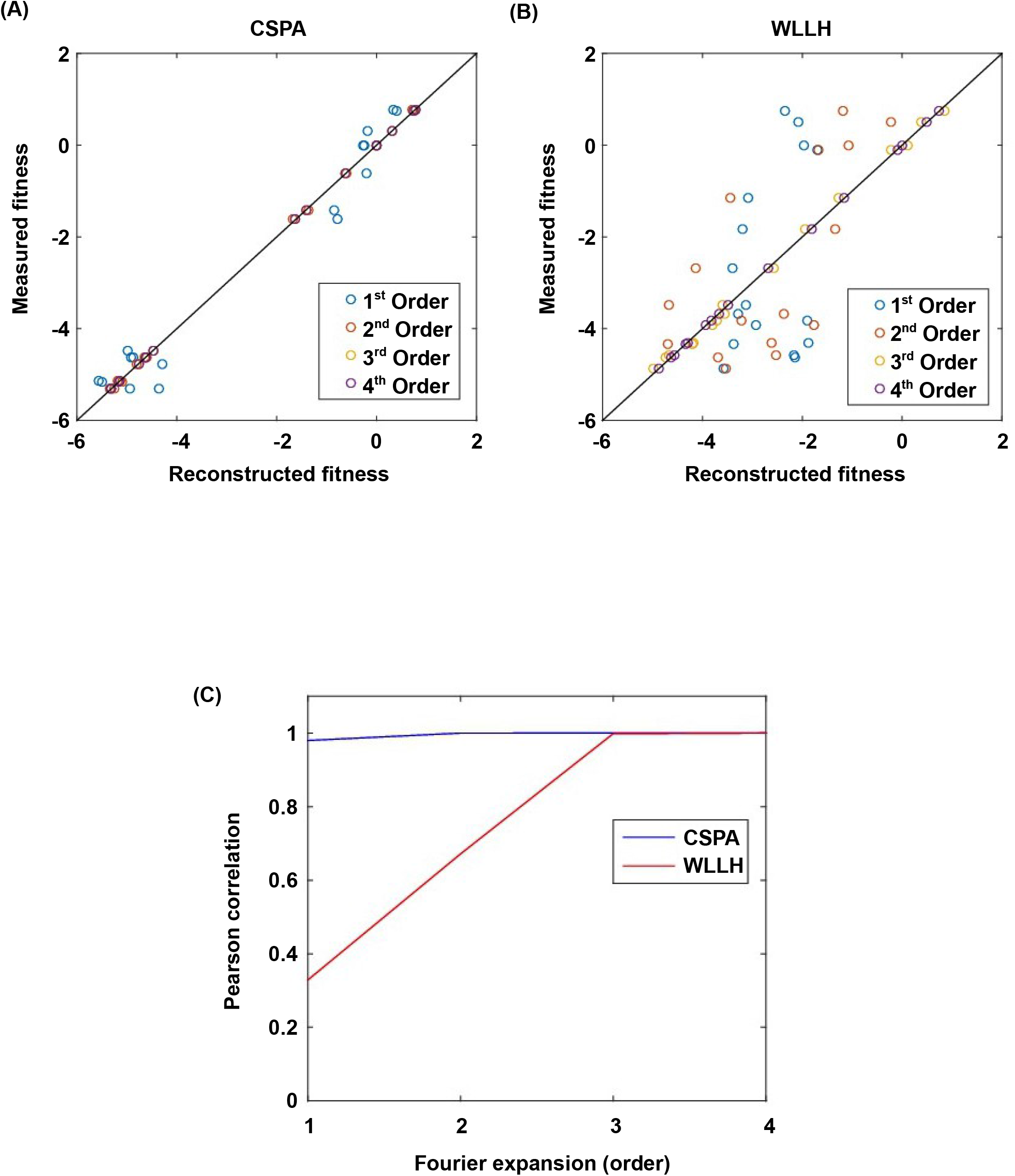
Fourier analysis decomposes the fitness landscape into epistatic interactions of different orders. Here we show two examples where the fitness contribution from higher-order epistasis is small (bottom 0.1%) in **(A)** and large (top 0.1%) in **(B)**. The fitness of variants can be reconstructed using Fourier coefficients truncated to a certain order. The Fourier coefficients can be interpreted as epistatic interactions of different orders, including the main effects of single mutations (the 1^*st*^ order), pair-wise epistasis (the 2^*nd*^ order), and higher-order epistasis (the 3^*rd*^ and the 4^*th*^ order). **(C)** In our system with four sites, the reconstructed fitness by expansion to the 4^*th*^ order Fourier coefficients will always reproduce the measured fitness (i.e. Pearson correlation equals 1). If expansion to the 2^*nd*^ order Fourier coefficients did not reproduce the measured fitness (i.e. Pearson correlation less than 1), it would indicate the presence of higher-order epistasis.

**Supplementary Figure 7.**
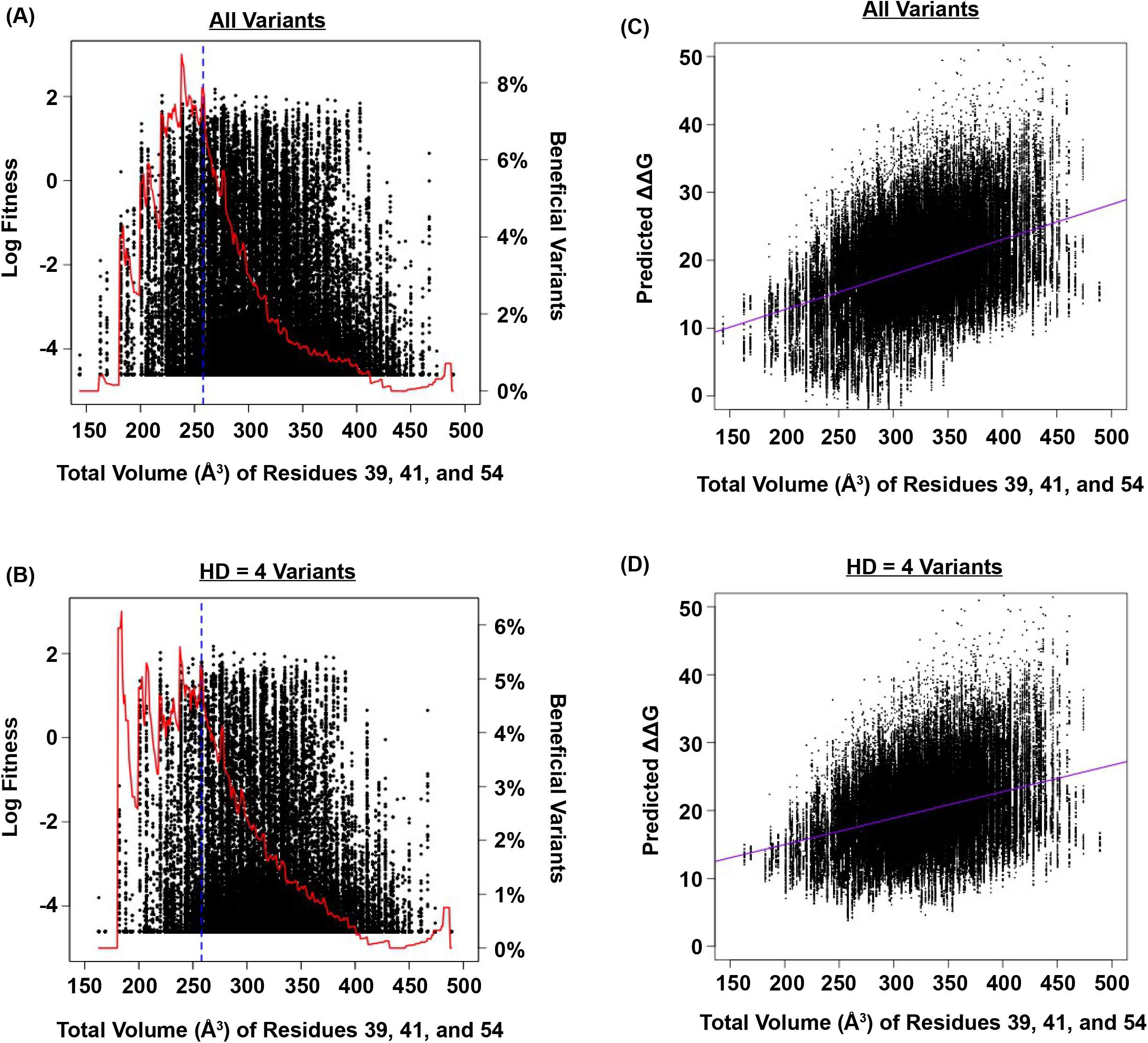
Relationship between fitness, size of the protein core, and predicted ∆∆G. **(A and B)** The relationship between fitness and the total volume of residue 39, 41, and 54 is shown as a scatter plot for **(A)** all variants, and **(B)** variants with HD of 4 from WT. The blue line indicates the volume of WT (VDGV). The red line indicates the fraction of beneficial variants within a sliding window of ±20 Å^3^. We postulated that the observed higher-order epistasis was, at least partially, due to the steric effect among site 39, 41, and 54. This was evidenced by the enrichment of beneficial variants when the total volume of these three interacting residues was between ∼200 Å^3^ and ∼300 Å^3^. As the total volume further increased, proportion of beneficial variants dropped. **(C and D)** The relationship between the predicted ∆∆G and the total size of residue 39, 41, and 54 for all variants is shown as a scatter plot for **(C)** all variants (Pearson’s correlation = 0.41), and **(D)** variants with HD of 4 from WT (Pearson’s correlation = 0.34). The purple line represents the linear regression. The predicted ∆∆G increased as the total volume of core residues increased, indicating that the protein would be destabilized (i.e. decrease in fitness) when the core was overpacked. Therefore, the higher-order epistasis observed in this study could be partially attributed to the steric effect. Nonetheless, we acknowledged that entropic effect and conformational effect in IgG-FC binding may also contribute to higher-order epistasis.

**Supplementary Figure 8.**
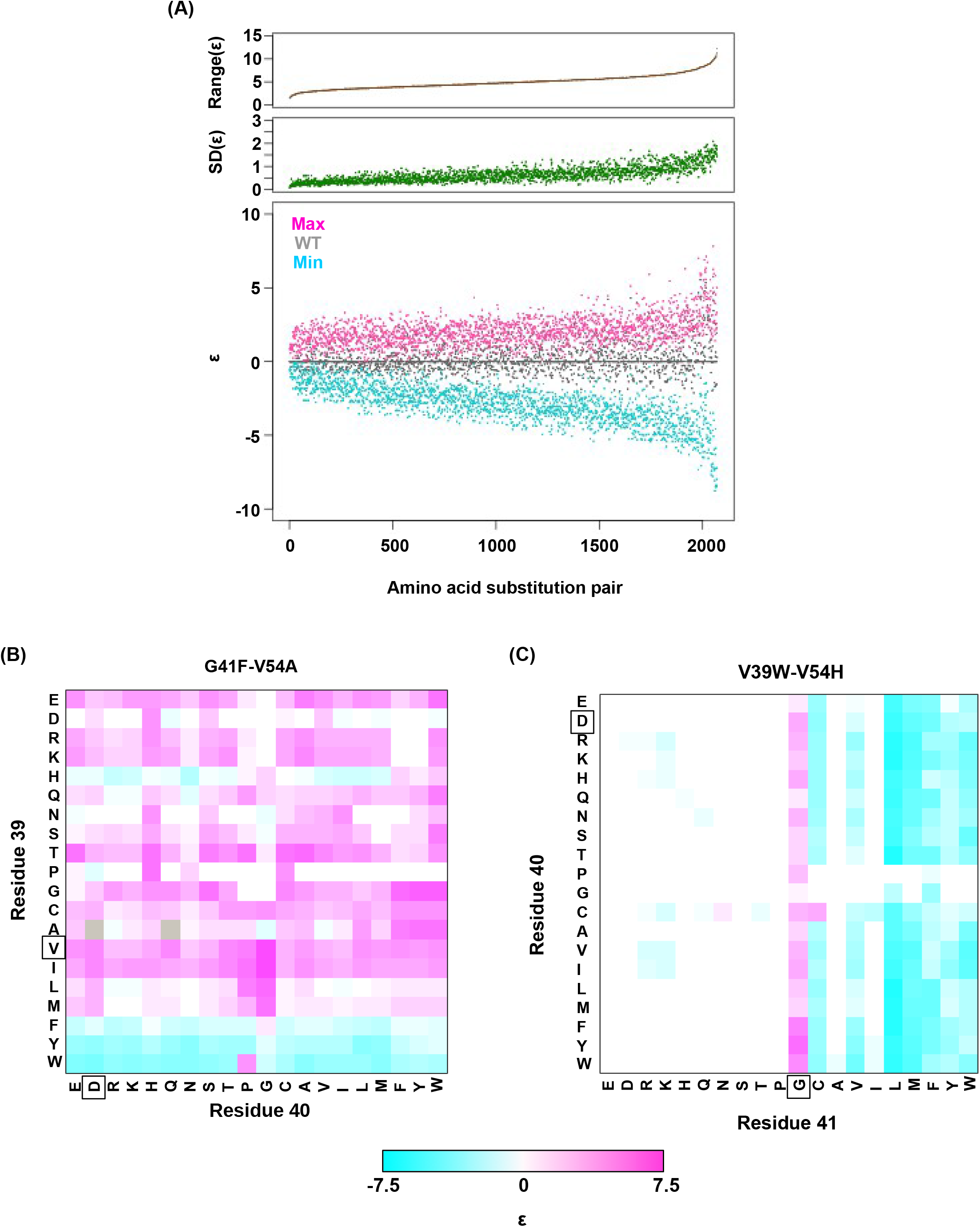
Alteration of pairwise epistatic effect under different genetic backgrounds. **(A)** Pairwise epistatic effect of each substitution pair under each genetic background (total possible genetic backgrounds for each substitution pair = 20 × 20 = 400) was quantified. For each substitution pair, the range of epistasis across different genetic backgrounds is shown in the top panel (brown). For each substitution pair, the standard deviation of epistasis across different genetic backgrounds is shown in the middle panel (green). For each substitution pair, the maximum epistatic value across different genetic backgrounds (ma-genta), the minimum epistatic value across different genetic backgrounds (cyan), and the epistatic value under WT background (grey) are shown. Substitution pair is ranked by the range of epistasis. **(B)** Epistasis between G41F and V54A across different genetic backgrounds (different combination of amino acids in sites 39 and 40) is shown. The epistasis value is color coded. Amino acids of WT are boxed. Epistasis that cannot be determined due to missing variant is colored in grey. **(C)** Epistasis between V39W and V54H across different genetic backgrounds (different combination of amino acids in sites 40 and 41) is shown. The epistasis value is color coded. Amino acids of WT are boxed. Epistasis that cannot be determined due to missing variant is colored in grey.

**Supplementary Figure 9.**
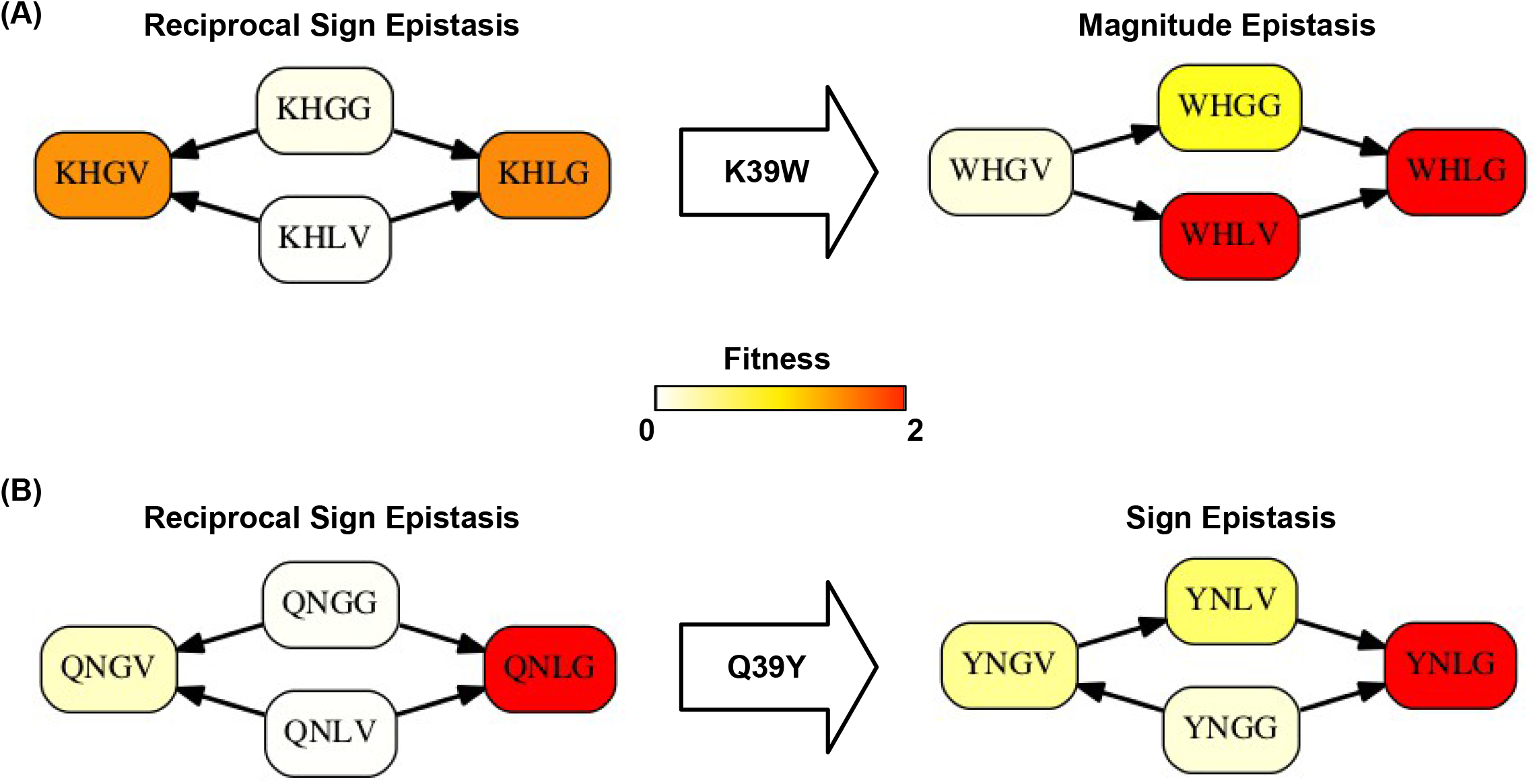
Higher-order epistasis can change the type of pairwise epistasis. The type of pairwise interaction could be changed in the presence of higher-order epistasis. **(A)** Reciprocal sign epistasis between G41L-V54G is changed to magnitude epistasis given the mutation at site 39 (K39W). **(B)** Reciprocal sign epistasis between G41L-V54G is changed to sign epistasis given the mutation at site 39 (Q39Y).

**Supplementary Figure 10.**
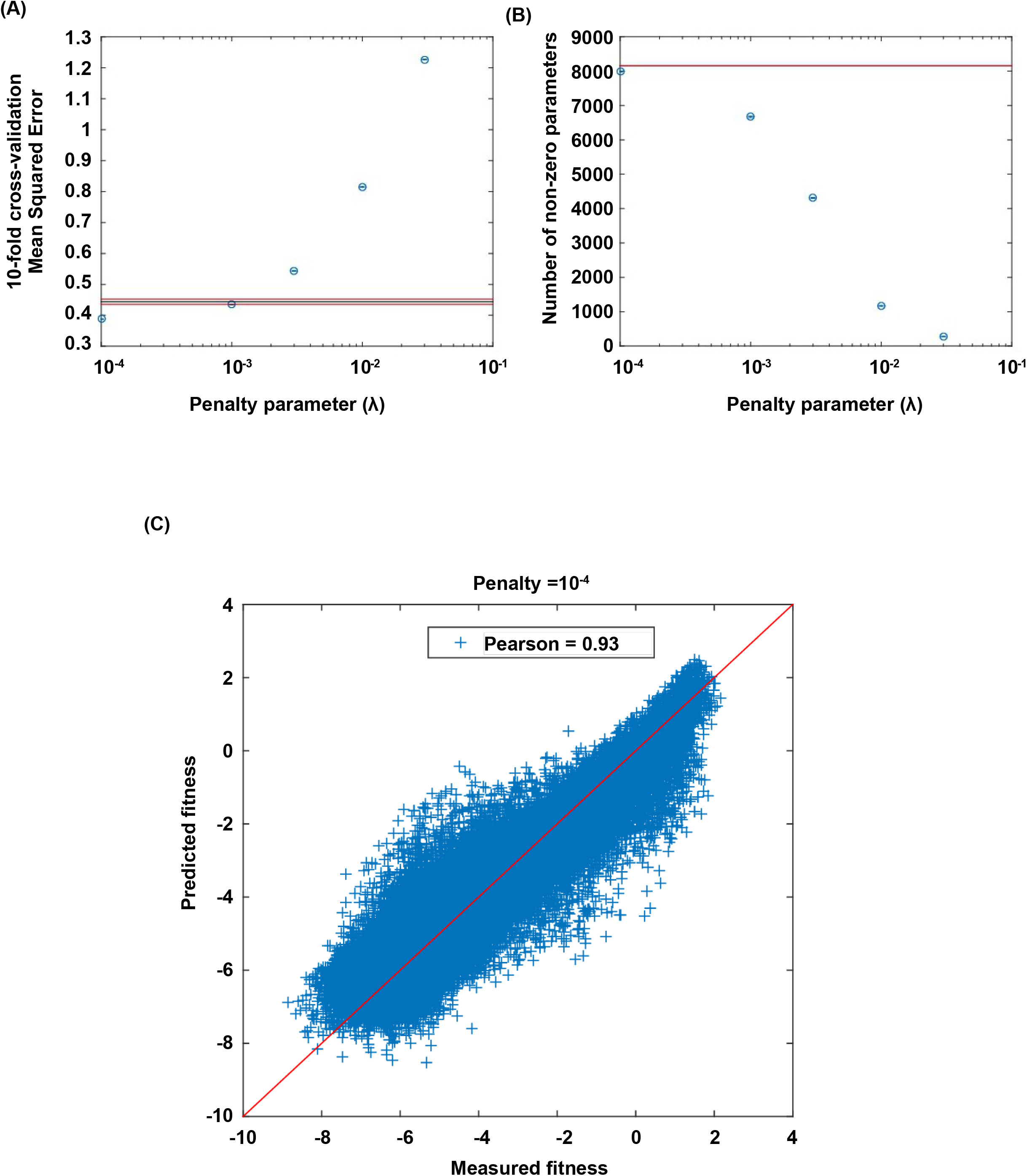
Lasso regression. Coefficients of the statistical model were fit by lasso regression on the measured fitness values of 119,884 non-lethal variants (see Methods). **(A)** 10-fold CV (cross-validation) MSE (mean squared errors) of lasso regression with varying penalty parameter λ. The black line indicates the 10-fold CV MSE of ordinary least squares regression (i.e. penalty parameter is zero). The red lines indicate the standard deviation. λ = 10^−4^ is chosen for imputing the fitness values of missing variants. **(B)** The number of nonzero coefficients in the model with varying penalty parameter λ. **(C)** Comparison between the predicted fitness values and the measured fitness values (Pearson correlation=0.93).

**Supplementary Figure 11.**
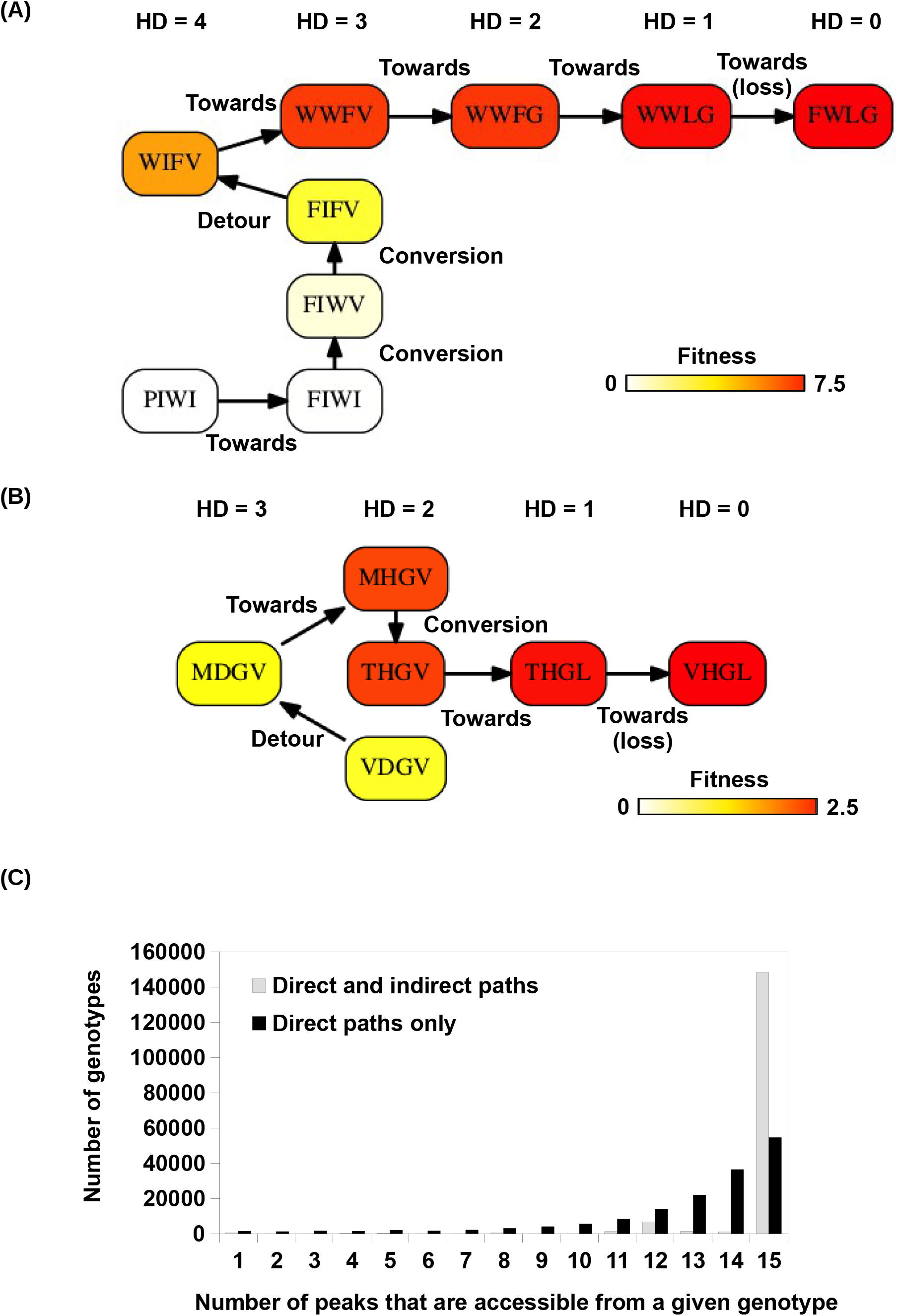
Indirect paths in adaptation. **(A)** A mutational trajectory initiated from PIWI under Greedy Model, which ended at the fitness peak, FWLG. **(B)** One of the shortest mutational trajectories from WT (VDGV) to a beneficial mutation (VHGL). **(C)** Histogram of the number of fitness accessible from a given genotype. The fraction of genotypes accessible to 15 fitness peaks increased substantially when indirect paths are allowed in adaptation.

**Supplementary Figure 12.**
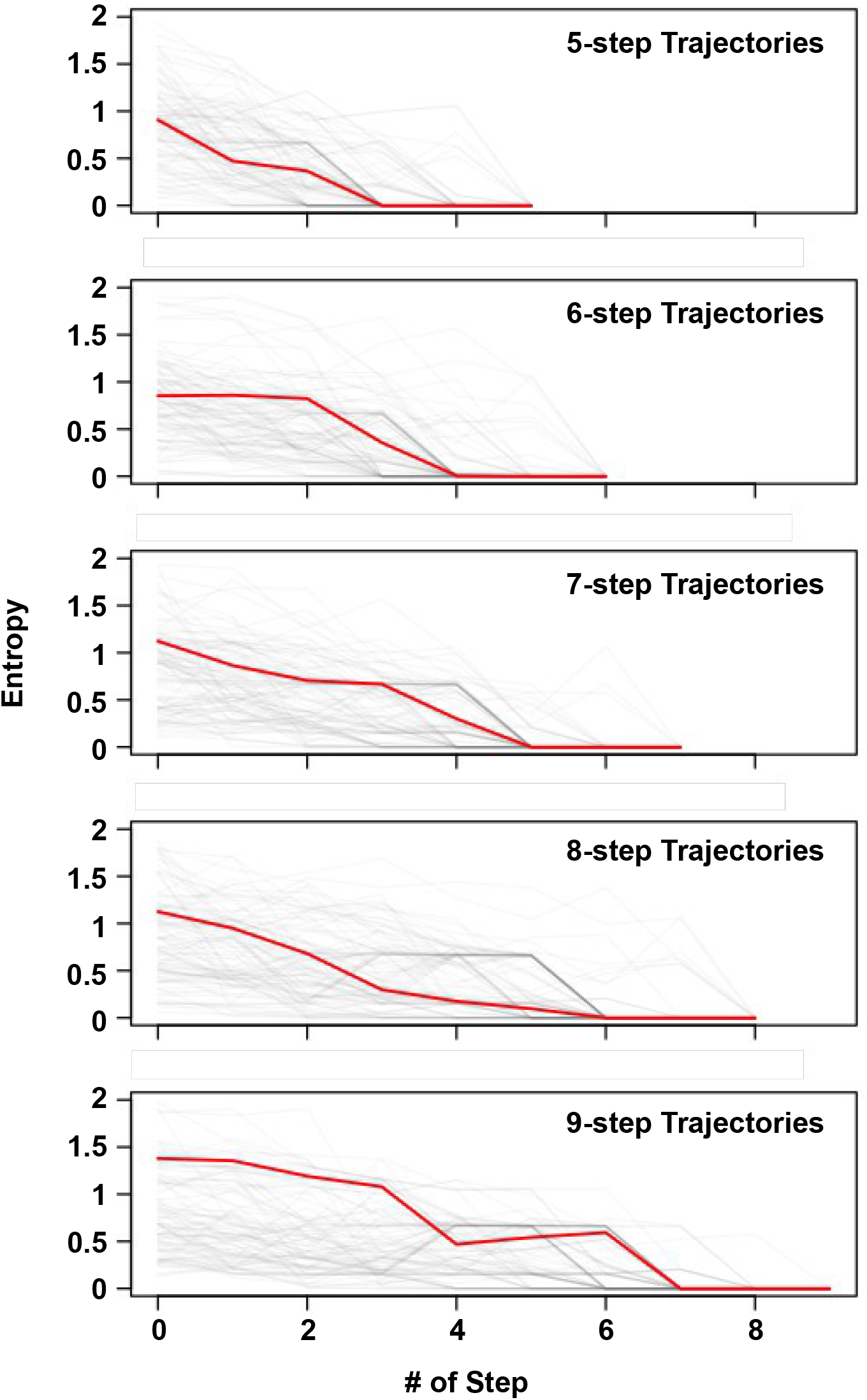
Delay of commitment in mutational trajectories involving extra-dimensional bypass. An entropy of evolutionary outcome was calculated for each of the 160,000 variants. Given a variant ***v*** with ***n*** accessible fitness peaks, the entropy of evolutionary outcome was then computed as follow:

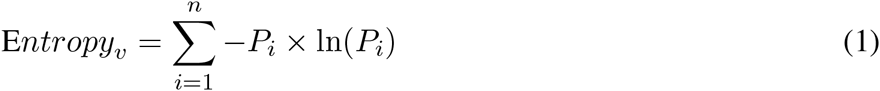

where *P_i_* represented the frequency of reaching the fitness peak *i* among 1,000 simulated mutational trajectories from variant *v* following Correlated Fixation Model. The entropy of evolutionary fates at each step along an adaptive path is shown. Adaptive paths with the same number of steps are grouped together. We observed that many mutational trajectories that involved extra-dimensional bypass did not fully commit to a fitness peak (entropy = 0) until the last two steps. Each grey line represents a mutational trajectory in each category. Only 100 randomly sampled trajectories are shown due to the difficulty in visualizing a large number of lines on the graph. The median entropy at each step in each category is represented by the red line.

## Supplementary Text

Here we prove that higher-order epistasis is required for two possible scenarios of extra-dimensional bypass via an additional site (Supplementary Fig. 5). For a fitness landscape defined on a Boolean hypercube, we can expand the fitness as Taylor series [51].

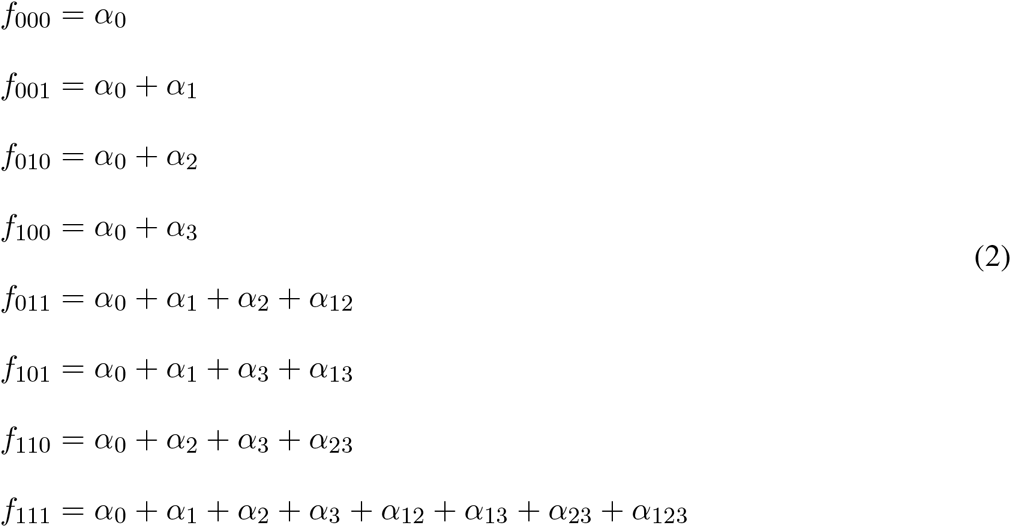

To prove that higher-order epistasis is present is equivalent to prove that α_123_ ≠ 0. The fitness difference between neighbors is visualized by the directed edges that go from low-fitness variant to high-fitness variant, thus each edge represents an inequality. No cyclic paths are allowed in this directed graph.

The reciprocal sign epistasis (Supplementary Fig. 5A) gives,

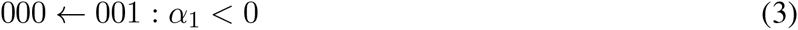

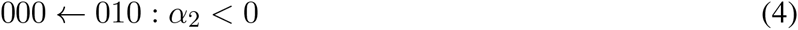

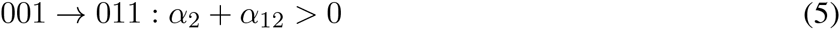

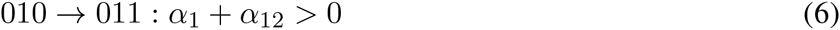

The detour step (000 → 100) and the loss step (111 → 011) are required for extra-dimensional bypass,

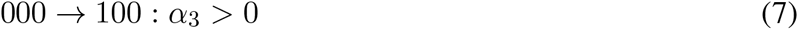

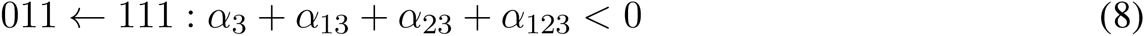

For the remaining 6 edges, there are 3 possible configurations (Supplementary Fig. 5B-D). For the scenario illustrated in **(B)**, we have

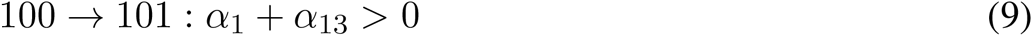

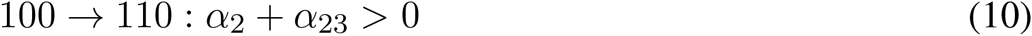

Combining inequality (3) and (9) gives

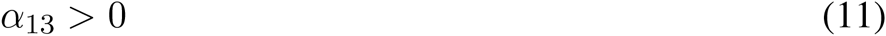

Combining inequality (4) and (10) gives

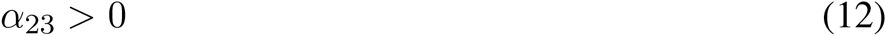

Combining the above two inequalities with (7) and (8), we arrive at

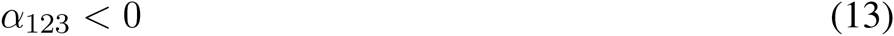

For the scenario in **(C)**, the proof of higher-order epistasis is similar. We have (the yellow edge)

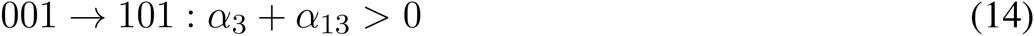

Combining the above inequality with (4), (8) and (10), we arrive at

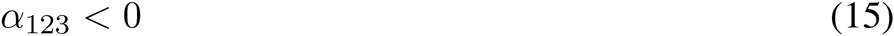

For the scenario in **(D)**, when α_3_ + α_13_ < 0, all the inequalities can be satisfied with α_123_ = 0. So higher-order epistasis is not necessary in this case.

## Methods

### Mutant library construction

Two oligonucleotides (Integrated DNA Technologies, Coralville, IA), 5’-AGT CTA GTA TCC AAC GGC NNS NNS NNK GAA TGG ACC TAC GAC GAC GCT ACC AAA ACC TT-3’ and 5’-TTG TAA TCG GAT CCT CCG GAT TCG GTM NNC GTG AAG GTT TTG GTA GCG TCG TCG T-3’ were annealed by heating to 95°C for 5 minutes and cooling to room temperature over 1 hour. The annealed nucleotide was extended in a reaction containing 0.5 uM of each oligonucleotide, 50 mM NaCl, 10 mM Tris-HCl pH 7.9, 10 mM MgCl_2_, 1 mM DTT, 250 uM each dNTP, and 50 units Klenow exo- (New England Biolabs, Ipswich, MA) for 30 mins at 37°C. The product (cassette I) was purified by PureLink PCR Purification Kit (Life Technologies, Carlsbad, CA) according to manufacturer’s instructions.

A constant region was generated by PCR amplification using KOD DNA polymerase (EMD Millipore, Billerica, MA) with 1.5 mM MgSO_4_, 0.2 mM of each dNTP (dATP, dCTP, dGTP, and dTTP), 0.05 ng protein GB1 wild type (WT) template, and 0.5 uM each of 5’-TTC TAA TAC GAC TCA CTA TAG GGA CAA TTA CTA TTT ACA TAT CCA CCA TG-3’ and 5’-AGT CTA GTA TCC TCG ACG CCG TTG TCG TTA GCG TAC TGC-3’. The sequence of the WT template consisted of a T7 promoter, 5’ UTR, the coding sequence of Protein GB1, 3’ poly-GS linkers, and a FLAG-tag (Supplementary Fig. 1B) [33]. The thermocycler was set as follows: 2 minutes at 95°C, then 18 three-step cycles of 20 seconds at 95°C, 15 seconds at 58°C, and 20 seconds at 68°C, and 1 minute final extension at 68°C. The product (constant region) was purified by PureLink PCR Purification Kit (Life Technologies) according to manufacturer’s in-structions. Both the purified constant region and cassette I were digested with BciVI (New England Biolabs) and purified by PureLink PCR Purification Kit (Life Technologies) according to manufacturer’s instructions.

Ligation between the constant region and cassette I (molar ratio of 1:1) was performed using T4 DNA ligase (New England Biolabs). Agarose gel electrophoresis was performed to separate the ligated product from the reactants. The ligated product was purified from the agarose gel using Zymoclean Gel DNA Re-covery Kit (Zymo Research, Irvine, CA) according to manufacturer’s instructions. PCR amplification was then performed using KOD DNA polymerase (EMD Millipore) with 1.5 mM MgSO_4_, 0.2 mM of each dNTP (dATP, dCTP, dGTP, and dTTP), 4 ng of the ligated product, and 0.5 uM each of 5’-TTC TAA TAC GAC TCA CTA TAG GGA CAA TTA CTA TTT ACA TAT CCA CCA TG-3’ and 5’-GGA GCC GCT ACC CTT ATC GTC GTC ATC CTT GTA ATC GGA TCC TCC GGA TTC-3’. The thermocycler was set as follows: 2 minutes at 95°C, then 10 three-step cycles of 20 seconds at 95°C, 15 seconds at 56°C, and 20 seconds at 68°C, and 1 minute final extension at 68°C. The product, which is referred as “DNA library”, was purified by PureLink PCR Purification Kit (Life Technologies) according to manufacturer’s instructions.

### Affinity selection by mRNA display

Affinity selection by mRNA display [34, 35] was performed as described (Supplementary Fig. 3A) [33]. Briefly, The DNA library was transcribed by T7 RNA polymerase (Life Technologies) according to man-ufacturer’s instructions. Ligation was performed using 1 nmol of mRNA, 1.1 nmol of 5’-TTT TTT TTT TTT GGA GCC GCT ACC-3’, and 1.2 nmol of 5-/5Phos/-d(A)21-(9)3-ACC-Puromycin by T4 DNA ligase (New England Biolabs) in a 100 uL reaction. The ligated product was purified by urea PAGE and translated in a 100 uL reaction volume using Retic Lysate IVT Kit (Life Technologies) according to manufacturer’s instructions followed by incubation with 500 mM final concentration of KCl and 60 mM final concentration of MgCl2 for at least 30 minutes at room temperature to increase the efficiency for fusion formation [52]. The mRNA-protein fusion was then purified using ANTI-FLAG M2 Affinity Gel (Sigma-Aldrich, St. Louis, MO). Elution was performed using 3X FLAG peptide (Sigma-Aldrich). The purified mRNA-protein fusion was reverse transcribed using SuperScript III Reverse Transcriptase (Life Technologies). This reverse tran-scribed product, which was referred as “input library”, was incubated with Pierce streptavidin agarose (SA) beads (Life Technologies) that were conjugated with biotinylated human IgG-FC (Rockland Immunochemicals, Limerick, PA). After washing, the immobilized mRNA-protein fusion was eluted by heating to 95°C. The eluted sample was referred as “selected library”.

### Sequencing library preparation

PCR amplification was performed using KOD DNA polymerase (EMD Millipore) with 1.5 mM MgSO_4_, 0.2 mM of each dNTP (dATP, dCTP, dGTP, and dTTP), the selected library, and 0.5 uM each of 5’-CTA CAC GAC GCT CTT CCG ATC T<u>NN N</u>AG CAG TAC GCT AAC GAC AAC G-3’ and 5’-TGC TGA ACC GCT CTT CCG ATC T<u>NN N</u>TA ATC GGA TCC TCC GGA TTC G-3’. The underlined “NNN” indicated the position of the multiplex identifier, GTG for input library and TGT for post-selection library. The thermocy-cler was set as follows: 2 minutes at 95°C, then 10 to 12 three-step cycles of 20 seconds at 95°C, 15 seconds at 56°C, and 20 seconds at 68°C, and 1 minute final extension at 68°C. The product was then PCR amplified again using KOD DNA polymerase (EMD Millipore) with 1.5 mM MgSO_4_, 0.2 mM of each dNTP (dATP, dCTP, dGTP, and dTTP), the eluted product from mRNA display, and 0.5 uM each of 5’-AAT GAT ACG GCG ACC ACC GAG ATC TA CAC TCT TTC CCT ACA CGA CGC TCT TCC G-3’ and 5’-CAA GCA GAA GAC GGC ATA CGA GAT CGG TCT CGG CAT TCC TGC TGA ACC GCT CTT CCG-3’. The thermocycler was set as follows: 2 minutes at 95°C, then 10 to 12 three-step cycles of 20 seconds at 95°C, 15 seconds at 56°C, and 20 seconds at 68°C, and 1 minute final extension at 68°C. The PCR product was then subjected to 2 × 100 bp paired-end sequencing using Illumina HiSeq 2500 platform. Raw sequencing data have been submitted to the NIH Short Read Archive under accession number: BioProject PRJNA278685.

We were able to compute the fitness for 93.4% of all variants from the sequencing data. The fitness measure-ments in this study were highly consistent with our previous study on fitness of single and double mutants in protein GB1 (Pearson correlation = 0.97, Supplementary Fig. 3B) [33].

### Sequencing data analysis

The first three nucleotides of both forward read and reverse read were used for demultiplexing. If the first three nucleotides of the forward read were different from that of the reverse read, the given paired-end read would be discarded. For both forward read and reverse read, the nucleotides that were corresponding to the codons of protein GB1 sites 39, 40, 41, and 54 were extracted. If coding sequence of sites 39, 40, 41, and 54 in the forward read and that in the reverse read did not reverse-complement each other, the paired-end read would be discarded. Subsequently, the occurrence of individual variants at the amino acid level for site 39, 40, 41, and 54 in both input library and selected library were counted, with each paired-end read represented 1 count. Custom python scripts and bash scripts were used for sequencing data processing. All scripts are available upon request.

### Calculation of fitness

The fitness (*w*) for a given variant *i* was computed as:

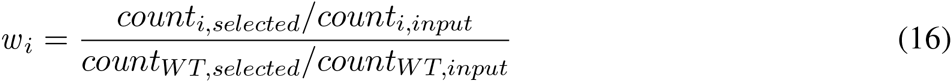

where count_*i,selected*_ represented the count of variant *i* in the selected library, count_*i,input*_ represented the count of variant i in the input library, count_*WT, selected*_ represented the count of WT (VDGV) in the selected library, and count_*WT, input*_ represented the count of WT (VDGV) in the input library.

Therefore, the fitness of each variant, *w_i_*, could be viewed as the fitness relative to WT (VDGV), such that *w_WT_* = 1. Variants with count_*input*_ < 10 were filtered to reduce noise. The fraction of all possible variants that passed this filter was 93.4% (149,361 out of 160,000 all possible variants).

The fitness of each single substitution variant was referenced to our previous study [33], because the se-quencing coverage of single substitution variants in our previous study was much higher than in this study (∼100 fold higher). Hence, our confidence in computing fitness for a single substitution variant should also be much higher in our previous study than this study. Subsequently, the fitness of each single substitution in this study was calculated by multiplying a factor of 1.159 by the fitness of that single substitution computed from our previous study [33]. This is based on the linear regression analysis between the single substitution fitness as measured in our previous study and in this study, which had a slope of 1.159 and a y-intercept of ∼0.

### Magnitude and type of pairwise epistasis

The three types of pairwise epistasis (magnitude, sign and reciprocal sign) were classified by ranking the fitness of the four variants involved [53].

To quantify the magnitude of epistasis (ε) between substitutions *a* and *b* on a given background variant *BG*, the relative epistasis model [39] was employed as follows:

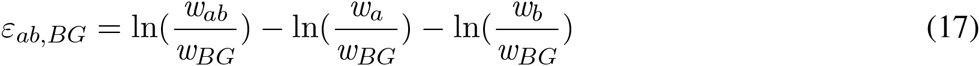

where *w_ab_* represents the fitness of the double substitution, ln(*w_a_*) and ln(*w_b_*) represents the fitness of each of the single substitution respectively, and *w_BG_* represents the fitness of the background variant.

As described previously [33], there exists a limitation in determining the exact fitness for very low-fitness variants in this system. To account for this limitation, several rules were adapted from our previous study to minimize potential artifacts in determining ε [33]. We previously determined that the detection limit of fitness (*w*) in this system is ∼0.01 [33].

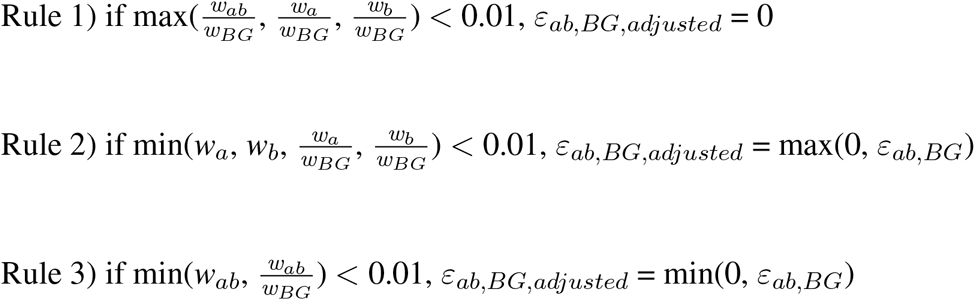

Rule 1 prevents epistasis being artifically estimated from low-fitness variants. Rule 2 prevents overestimation of epistasis due to low fitness of one of the two single substitutions. Rule 3 prevents underestimation of epistasis due to low fitness of the double substitution. To compute the epistasis between two substitutions, *a* and *b*, on a given background variant *BG*, *ε_ab, BG, adjusted_* would be used if one of the above three rules was satisfied. Otherwise, *ε_ab, BG_* would be used.

### Fourier analysis

Fitness decomposition was performed on all subgraphs without missing variants (109,235 subgraphs in total). We decomposed the fitness landscape into epistatic interactions of different orders by Fourier analysis [9, 54]. The Fourier coefficients given by the transform can be interpreted as epistasis of different orders [6,30].

For a binary sequence 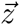 with dimension *L* (*z_i_* equals 1 if mutation is present at position *i*, or 0 otherwise), the Fourier decomposition theorem states that the fitness function 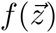 can be expressed as [51]:

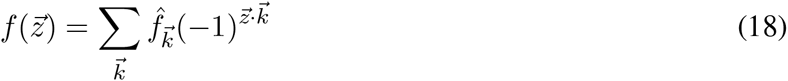

The formula for the Fourier coefficients 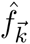 is then:

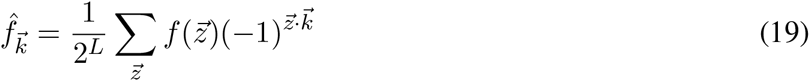

For example, we can expand the fitness landscape up to the second order, i.e. with linear and quadratic terms

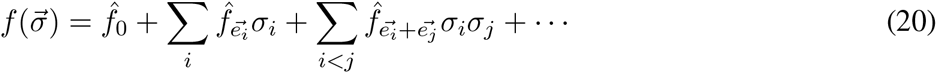

where 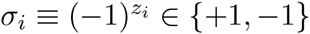, and 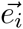 is a unit vector along the *i^th^* direction. In our analysis of subgraphs, there are a total of 2^4^ = 16 terms in the Fourier decomposition, with 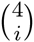 terms for the *i^th^* order (*i* = 0, 1, 2, 3, 4). We can expand the fitness landscape up to a given order by ignoring all higher-order terms in Equation 18. In this paper, we refer to higher-order epistasis as non-zero contribution to fitness from the 3^*rd*^ order terms and beyond.

### Imputing the fitness of missing variants

The fitness values for 10,639 variants (6.6% of the entire sequence space) were not directly measured (read count in the input pool = 0) or were filtered out because of low read counts in the input pool (see section “Calculation of fitness”). To impute the fitness of these missing variants, we performed regularized regression on fitness values of observed variants using the following model [40,55]:

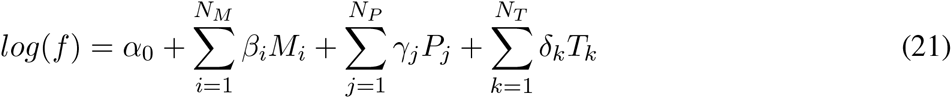

Here, *f* is the protein fitness. α_0_ is the intercept that represents the log fitness of WT; *β_i_* represents the main effect of a single mutation, *i*; *M_i_* is a dummy variable that equals 1 if the single mutation *i* is present in the sequence, or 0 if the single mutation is absent; and 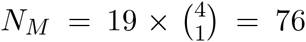 is the total number of single mutations. Similarly, *γ_j_* represents the effect of interaction between a pair of mutations; *P_j_* is the dummy variable that equals either 1 or 0 depending on the presence of that those two mutations; and 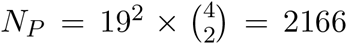 is the total number of possible pairwise interactions. In addition to the main effects of single mutations and pairwise interactions, the three-way interactions among sites 39, 41 and 54 are included in the model, based on our knowledge of higher-order epistasis (Fig. 3). *δ_k_* represents the effect of three-way interactions among sites 39, 41 and 54; *T_k_* is the dummy variable that equals either 1 or 0 depending on the presence of that three-way interaction; and *N_T_* = 19^3^ = 6859 is the total number of three-way interactions. Thus, the total number of coefficients in this model is 9,102, including main effects of each site (i.e. additive effects), interactions between pairs of sites (i.e. pairwise epistasis), and a subset of three-way interactions (i.e. higher-order epistasis).

Out of the 149,361 variants with experimentally measured fitness values, 119,884 variants were non-lethal (*f* > 0) and were used to fit the model coefficients using lasso regression (Matlab R2014b). Lasso regression adds a penalty term 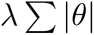 (*θ* stands for any coefficient in the model) when minimizing the least squares, thus it favors sparse solutions of coefficients (Supplementary Fig. 10B). We calculated the 10-fold cross-validation MSE (mean squared errors) of the lasso regression for a wide range of penalty parameter λ (Supplementary Fig. 10A). λ = 10^−4^ is chosen. For measured variants, the model-predicted fitness values were highly correlated with the actual fitness values (Pearson correlation=0.93, Supplementary Fig. 10C). We then used the fitted model to impute the fitness of the 10,639 missing variants and complete the entire fitness landscape.

### Simulating adaptation using three models for fixation

Python package “networkx” was employed to construct a directed graph that represented the entire fitness landscape for sites 39, 40, 41, and 54. A total of 4^20^ = 160,000 nodes were present in the directed graph, where each node represented a 4-site variant. For all pairs of variants separated by a Hamming distance of 1, a directed edge was generated from the variant with a lower fitness to the variant with a higher fitness. Therefore, all successors of a given node had a higher fitness than the given node. A fitness peak was defined as a node that had 0 out-degree. Three models, namely the Greedy Model [6], the Correlated Fixation Model [41], and the Equal Fixation Model [20], were employed in this study to simulate the mutational steps in adaptive trajectories. The Greedy Model represents adaptive evolution of a large population with pervasive clonal interference [6]. The Correlated Fixation Model represents adaptive evolution of a population under the scheme of strong-selection/weak-mutation (SSWM), which assumes that the time to fixation is much shorter than the time between mutations, and the fixation probability of a given mutation is proportional to the improvement in fitness. The Equal Fixation Model represents a simplified scenario of adaptation where all beneficial mutations fix with equal probability [20]. Under all three models, the probability of fixation of a deleterious or neutral mutation is 0. Considering a mutational trajectory initiated at a node, n_*i*_ with a fitness value of *w_i_*, where n_*i*_ has M successors, (n_1_, n_2_, … n_M_) with fitness values of (*w*_1_, *w*_2_, … *w*_M_). Then the probability that the next mutational step is from n_*i*_ to n_*k*_, where *k* ∈ (1, 2, … M), is denoted P_*i*→*k*_ and called the probability of fixation, and can be computed for each model as follows.

For the Greedy Model (deterministic model),

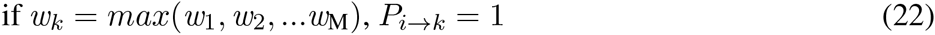

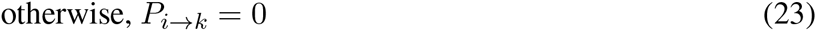

For the Correlated Fixation Model (non-deterministic model),

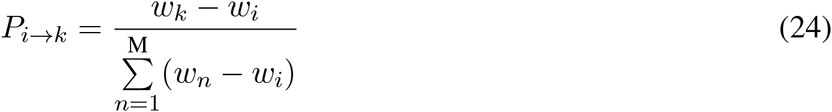

For the Equal Fixation Model (non-deterministic model),

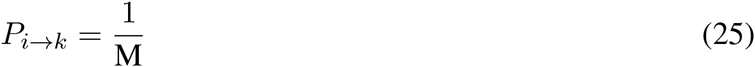

To compute the shortest path from a given variant to all reachable variants, the function “single_source_shortest_path” in “networkx” was used. If the shortest path between a low-fitness variant and a high-fitness variant does not exist, it means that the high-fitness variant is inaccessible. If the shortest path is longer than the Hamming Distance between two variants, it means that adaptation requires indirect paths.

### Analysis of direct paths within a subgraph

In the subgraph analysis shown in Supplementary Fig. 4, the fitness landscape was restricted to 2 amino acids at each of the 4 sites (the WT and adapted alleles). There was a total of 2^4^ variants, hence nodes, in a given subgraph. Only those subgraphs where the fitness of all variants was measured directly were used (i.e. any subgraph with missing variants was excluded from this analysis). Mutational trajectories were generated in the same manner as in the analysis of the entire fitness landscape (see subsection “Simulating adaptation using three models for fixation”). In a subgraph with only one fitness peak, the probability of a mutational trajectory from node *i* to node *j* via intermediate *a*, *b*, and *c* was as follows:

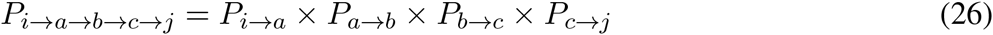

To compute the Gini index for a given set of mutational trajectories from node i to node j, the probabilities of all possible mutational trajectories were sorted from large to small. Inaccessible trajectories were also included in this sorted list with a probability of 0. This sorted list with *t* trajectories was denoted as 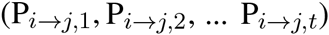, where 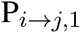 was the largest and 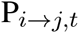 was the smallest. This sorted list was converted into a list of cumulative probabilities, which is denoted as 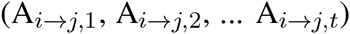, where 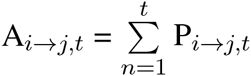

The Gini index for the given subgraph was then computed as follows:

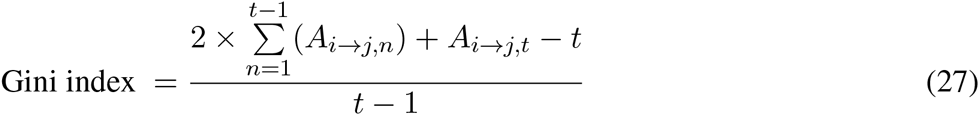

### Visualization

Sequence logo was generated by WebLogo (http://weblogo.berkeley.edu/logo.cgi) [56].

The visualization of basins of attraction (Fig. 4A) was generated using Graphviz with “fdp” as the option for layout.

### ∆∆G prediction

The ∆∆G prediction was performed by the ddg monomer application in Rosetta software [57] with the parameters from row 16 of Table 1 in Kellogg et al. were used [58].

## Acknowledgments

We would like to thank Jesse Bloom and Joshua Plotkin for helpful comments on early versions of the manuscript. N.C.W. was supported by Philip Whitcome Pre-Doctoral Fellowship, Audree Fowler Fellow-ship in Protein Science, and UCLA Dissertation Year Fellowship. L.D. was supported by HHMI Postdoc-toral Fellowship from Jane Coffin Childs Memorial Fund for Medical Research. R.S. was supported by NIH R01 DE023591. The funders had no role in study design, data collection and analysis, decision to publish, or preparation of the manuscript.

## Contributions

N.C.W., C.A.O., and R.S. designed the experiment, N.C.W. and C.A.O. performed the experiments, N.C.W. processed the sequencing data, L.D. and N.C.W. analyzed the fitness landscape, J.O.L.S. provided important intellectual inputs, L.D. and N.C.W. wrote the manuscript, J.O.L.S. and R.S. revised the manuscript.

